# 5-azacytosine induces cytotoxicity via 5-methylcytosine depletion on chromatin-associated RNA in leukemia

**DOI:** 10.1101/2025.06.24.661348

**Authors:** Boyang Gao, Ying Li, Long Zhao, Zhongyu Zou, Xinyuan Ma, Juyeong Hong, Junhong Xiang, Xiaoyang Dou, Feng-Chun Yang, Mingjiang Xu, Chuan He

## Abstract

5-azacytidine (5-azaC) is a DNA hypomethylating agent clinically used to improve outcomes in myeloid malignancies. However, 5-azaC treatment causes gene dysregulation inconsistent with DNA hypomethylation changes, suggesting alternative mechanisms of action by 5-azaC. As a ribonucleoside analogue, 5-azaC is more readily incorporated into nascent RNA. Here, we demonstrate that RNA 5-methylcytosine (m^5^C) depletion by 5-azaC treatment, particularly at early time points, is sufficient to induce leukemia cell death. In contrast to its DNA demethylation function, the RNA-dependent effect of 5-azaC causes transcriptional repression, disrupting genes involved in cell cycle regulation and DNA repair. Mechanistically, 5-azaC impairs two specific m^5^C-mediated transcriptional regulatory pathways. First, depletion of m^5^C in chromatin-associated RNA (caRNA) disrupts the MBD6-mediated H2AK119ub deubiquitination. In parallel, this also impairs SRSF2 recruitment and the downstream H3K27ac deposition by p300. Indeed, loss of the caRNA methyltransferase NSUN2 caused prolonged cell cycle, defective DNA repair, and shifted hematopoietic lineage commitment toward erythropoiesis, mirroring the effects of 5-azaC treatment. Furthermore, we performed a leukemia cell line screen and identified that TET2 and IKZF1 depletion can sensitize 5-azaC treatment, consistent with the observed RNA-dependent cytotoxicity of 5-azaC in leukemic cells. In summary, our findings highlight the transcription repression by 5-azaC through depleting caRNA m^5^C, providing additional insights into the mechanism of action for 5-azaC, the prediction of its efficacy, and future directions for therapy developments based on 5-azaC.

**HIGHLIGHT:**

- RNA-dependent effects of 5-azaC are sufficient to drive leukemia cell cytotoxicity through transcriptional repression.
- 5-azaC-induced caRNA m^5^C depletion impairs MBD6 binding and H2AK119ub deubiquitination.
- 5-azaC-induced caRNA m^5^C depletion disrupts SRSF2 chromatin-binding, impeding p300 recruitment and H3K27ac deposition.
- TET2 or IKZF1 depletion synergizes leukemia sensitivity to 5-azaC

## INTRODUCTION

5-azaC is the first-line therapy for high-risk myelodysplastic syndromes (MDS)^1^ and acute myeloid leukemia (AML) in older adults not eligible for intensive therapies^2^. The response rate of 5-azaC varies around 50%^3,4^, with frequent resistance development that limits its efficacy. Currently, there are no reliable biomarkers of 5-azaC efficacy prior to administration, indicating a lack of full understanding of its mechanism of action.

As a ribonucleoside analog of cytidine, 5-azaC can be incorporated into nucleic acids, with 10%-20% integrated into DNA and 80%-90% into RNA^5^. Interestingly, previous mechanistic investigations predominantly focused on the DNA-dependent cytotoxicity of 5-azaC, highlighting its role as a hypomethylation agent that covalently inhibits DNA methyltransferases^6–8^, thereby disrupting DNA 5mC deposition and epigenetic regulation. However, DNA demethylation targets do not fully align with the differentially expressed genes following 5-azaC treatment^9–12^, especially the transcription repression effect of 5-azaC, suggesting alternative mechanisms of action. In addition to DNA methyltransferases, 5-azaC also impairs the activity of RNA m^5^C methyltransferases^13,14^, whose effects and underlying pathways on leukemia cell cytotoxicity have been largely overlooked. The deoxy form of 5-azaC, decitabine, which exclusively incorporates into DNA, exhibits notably weaker effects on MDS and AML treatment^15–18^. This suggests a significant contribution of the RNA-dependent cytotoxicity for 5-azaC^19^. Additionally, patients benefited from 5-azaC administration did not necessarily exhibit DNA hypomethylation^20^. Moreover, it has been indicated that sensitivity to 5-azaC can be associated with expression levels and regulatory roles of RNA m^5^C methyltransferases^21^. These findings altogether indicate a crucial role of RNA m^5^C depletion in 5-azaC treatment, yet detailed mechanisms connecting RNA m^5^C depletion with cytotoxicity of 5-azaC are still lacking.

RNA m^5^C regulates multiple cellular processes^22,23^. In rRNA, m^5^C modifications influence ribosomal biogenesis^24^, while in tRNA, they impact RNA stability^25^, both contributing to translational regulation. In mRNA, m^5^C may affect splicing^26^, nuclear export^27^, and RNA stability^28,29^, likely mediated by the m^5^C-binding proteins SRSF2, ALYREF, or YBX1, respectively. We have recently discovered a chromatin and transcriptional regulatory role of m^5^C in chromatin-associated RNA (caRNA)^30^. We found that caRNA m^5^C can be recognized by MBD6^30^ and MBD5^31^, which then recruit the BAP1 complex (PR-DUB) for the removal of histone H2AK119ub to confer transcriptional activation. Oxidation of caRNA m^5^C by the TET2 enzyme suppresses this gene activation pathway^30^.

In this study, we characterized the RNA-dependent cytotoxicity of 5-azaC by monitoring its cellular responses at early time points. Our results show that the RNA-dependent pathways were sufficient to induce leukemia cell death, disrupting cell cycle regulation and DNA damage repair, and promoting apoptosis. Contrary to the conventional view of 5-azaC as a DNA hypomethylating agent, its RNA-dependent effects caused transcriptional repression, contributing to the cellular dysregulation. Mechanistically, 5-azaC depletes m^5^C on chromatin-associated repeat RNA, reducing MBD6 binding and subsequent recruitment of the BAP1 deubiquitylase complex for histone H2AK119ub removal. In parallel, caRNA m^5^C depletion also disrupts SRSF2 binding to chromatin, impairing p300 recruitment and H3K27ac deposition. These transcriptional effects by 5-azaC are primarily mediated through an inhibition of NSUN2, an RNA methyltransferase responsible for caRNA m^5^C deposition. Indeed, NSUN2 depletion recapitulated the cellular dysregulation observed with 5-azaC treatment at early time points. We further showed that *Nsun2* KO in mice led to shifted hematopoietic commitment towards the erythroid lineage, similar to what was observed in 5-azaC-treated patients. Finally, our leukemia cell line screen identified TET2 and IKZF1 mutations as indicators of 5-azaC sensitivity. These findings highlight the significance of the RNA-dependent pathways in 5-azaC treatment, and provide new insights into efficacy prediction and novel therapy development based on 5-azaC.

## RESULTS

### 5-azaC induces RNA m^5^C depletion and leukemia cell death at early time points

We reasoned that 5-azaC could be readily incorporated into nascent RNA, whereas its incorporation into DNA requires prior conversion to its deoxy form. RNA-mediated effects of 5-azaC are likely most evident at early treatment stages, before substantial incorporation into DNA occurs^19^. We first measured RNA m^5^C and DNA 5mC levels using ultra-high-performance liquid chromatography-tandem mass spectrometry (UHPLC-MS/MS), given that 5-azaC could covalently inhibit both RNA and DNA cytosine methyltransferases. 5-azaC was administered at 3 μM, a concentration relevant to its plasma levels (∼11 μM) in standard treatment^32^. Both tRNA and caRNA exhibited significantly decreased m^5^C levels starting at 8 hours, with caRNA showing a greater decrease (Figure 1A). In contrast, 5mC levels in genomic DNA were not significantly changed within 24 hours (Figure 1A), supporting our speculation that the early effects of 5-azaC treatment should largely arise from RNA m^5^C changes.

**Figure 1.**
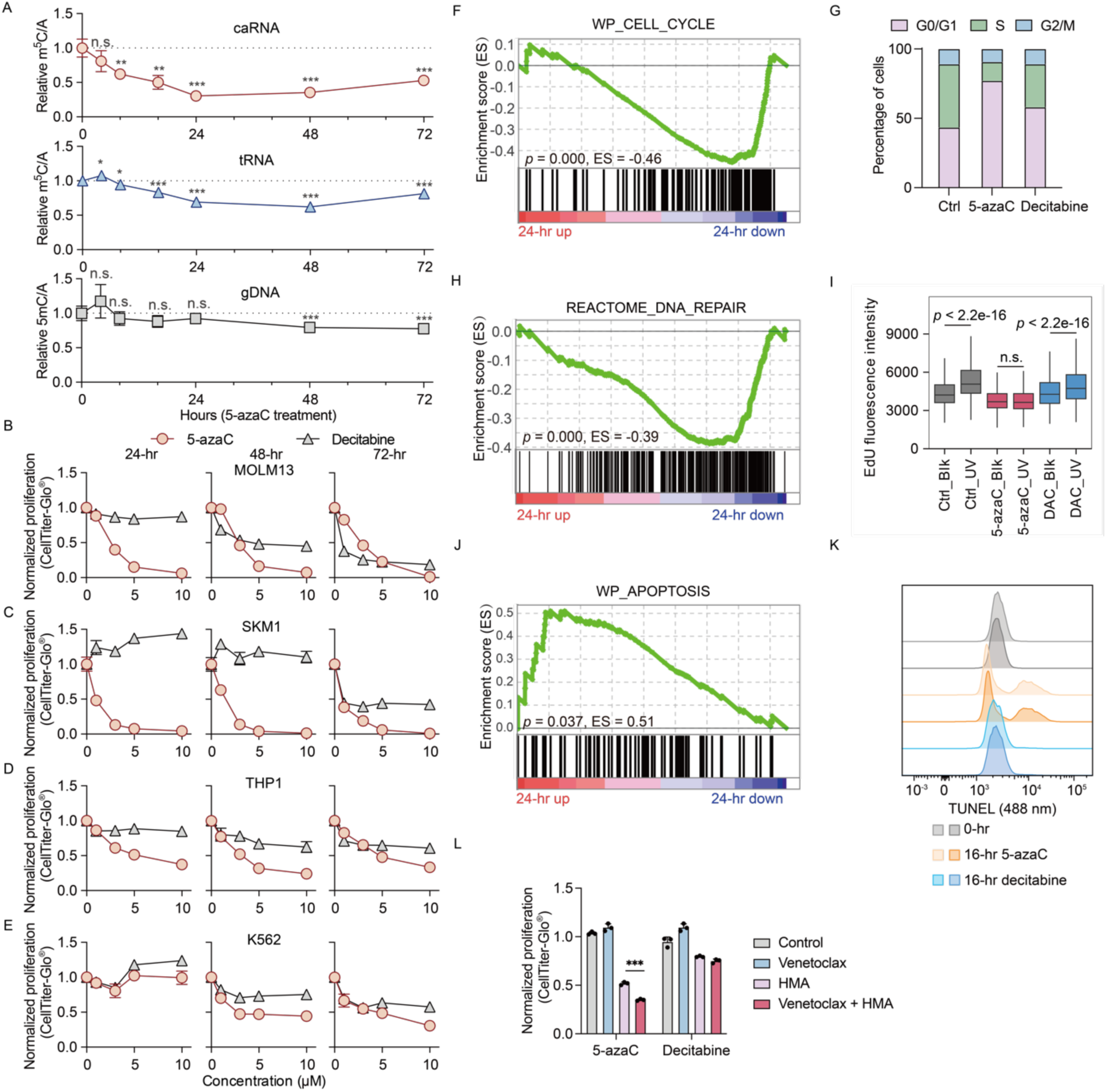
5-azaC induces RNA m^5^C depletion and leukemia cell death at early time points. (A) Changes of m^5^C/A levels in caRNA and tRNA, and 5mC/A levels in gDNA after 5-azaC (3 μM) treatment in MOLM13. Modification levels are normalized to controls. (B-E) Cell proliferation of MOLM13 (B), SKM-1 (C), THP1 (D) and K562 (E) after 5-azaC or decitabine treatment. (F) GSEA of MSigDB cell cycle gene set in 5-azaC-treated MOLM13. (G) Cell cycle distribution of MOLM13 cells after a 16-hour 5-azaC or decitabine treatment. (H) GSEA of MSigDB DNA repair gene set in 5-azaC-treated MOLM13. (I) EdU incorporation during DNA damage repair after UV irradiation in MOLM13 cells treated with 3 μM of 5-azaC or decitabine for 16 hours. Only G1-phase populations are shown. Blk, no irradiation. DAC, decitabine. (J) GSEA of MSigDB apoptosis gene set in 5-azaC-treated MOLM13 cells. (K) TUNEL fluorescent signals in MOLM13 cells following 3 μM of 5-azaC or decitabine treatment. (L) Cell proliferation of MOLM13 after 3 μM of 5-azaC or decitabine treatment in combination with 0.2 nM venetoclax for 24 hours. HMA, hypomethylating agent.

Next, we performed a proliferation inhibition assay using different leukemia cell lines. Within the clinically relevant concentration range^32^ of ∼10 μM, proliferation assays showed that 5-azaC induces significant cell death within 24 hours in different leukemia cell lines; whereas its deoxy form, decitabine, which could only be incorporated into DNA, had little impact within the same treatment window (Figure 1B-E). These findings support that the RNA-dependent pathways alone can cause leukemia cell death. Sensitivity to 5-azaC varied across leukemia cell lines: MOLM13 and SKM1, both derived from AML patients with prior MDS progression, were more susceptible to 5-azaC treatment at early time points (Figure 1B and C). In contrast, the acute monocytic leukemia cell line THP1 displayed lower sensitivity (Figure 1D), and K562 showed no response within 24 hours (Figure 1E).

We sought to further understand the RNA-dependent pathways underlying the cytotoxicity of 5-azaC. RNA-seq analysis following 24-hour 5-azaC treatment in MOLM13 and SKM1 cells identified differentially expressed genes (DEGs), with both gene sets enriched in pathways associated with cancer cell survival. Gene ontology (GO) analysis of biological processes showed that downregulated DEGs were involved in cell division, DNA repair, and RNA processing, while upregulated DEGs were linked to the inflammatory response (Figure S1A-D).

We first focused on cell division to explain the reduced cell proliferation upon 5-azaC treatment. Multiple pathways related to cell division were downregulated, including mitotic cell cycle, DNA replication, and chromosome segregation (Figure S1C and D). Gene set enrichment analysis (GSEA) revealed downregulation of cell cycle-related genes (Figure 1F). A 16-hour treatment with 5-azaC significantly decreased the S-phase cell population in MOLM13 and SKM1, whereas decitabine treatment only caused a minor reduction in MOLM13 proliferation (Figure 1G and S1E). These observations suggest that 5-azaC treatment at early time points prolongs the leukemia cell cycle, leading to proliferation inhibition.

Additionally, GO analysis revealed downregulation of DNA damage repair genes 24 hours after 5-azaC treatment (Figure S1C and D). GSEA showed a collective downregulation of DNA repair pathway genes (Figure 1H). The downregulation was further confirmed by immunoblotting of several prominently changed genes, including those encoding central checkpoint kinase CHK1 and DNA repair protein RAD51 (Figure S1F). Notably, UVC irradiation elicited diminished γH2A.X signaling following 5-azaC treatment in MOLM13 cells (Figure S1G). UVC-induced unscheduled DNA synthesis (UDS), assessed via 5-ethynyl-2’-deoxyuridine (EdU) incorporation and click chemistry, demonstrated impaired DNA repair activity after a 16-hour exposure to 5-azaC, but not to decitabine (Figure 1I and S1H). These findings further uncover that the early effects of 5-azaC also disrupt DNA damage response in leukemic cells. The accumulation of DNA lesions may further impede cell cycle progression, enhancing the prolonged cell cycle observed with 5-azaC treatment (Figure 1G and S1E).

As previously established, defective cell cycle progression induces apoptosis. In parallel, apoptosis also arises from unsuccessful DNA repair, and DNA damage response inhibitors have been used in cancer therapy to enhance cell death^33^. The defective cell cycle regulation and DNA repair may explain the cytotoxic effects of 5-azaC by inducing apoptosis. Consistently, GSEA showed upregulation of apoptotic genes (Figure 1J). The terminal deoxynucleotidyl transferase dUTP nick end labeling (TUNEL) assay detected apoptosis in MOLM13 cells after a 16-hour treatment with 5-azaC but not decitabine (Figure 1K). Intriguingly, 5-azaC has been combined with the BCL2 inhibitor venetoclax in clinical usage, resulting in improved recovery in AML patients^34,35^. Indeed, when we combined 5-azaC and venetoclax, we observed enhanced efficacy of proliferation inhibition for MOLM13 cells after 24 hours, whereas no synergistic effect was observed for decitabine and venetoclax under the same conditions (Figure 1L). These results support RNA-dependent cytotoxicity of 5-azaC in leukemic cells at early time points, and this effect can synergize with pro-apoptotic agents.

To further explore the RNA-dependent effect of 5-azaC under physiological conditions, we performed a 24-hour 5-azaC treatment of the Lin^−^KIT^+^ (LK) cells purified from mouse bone marrow (BM). GSEA analysis revealed downregulation of cell cycle and DNA repair genes similar to those observed in leukemic cells after the same treatment (Figure S1 I-J). Our findings therefore indicate cytotoxicity of 5-azaC mediated through RNA m^5^C depletion at early time points.

### 5-azaC induces transcriptional repression through caRNA m^5^C depletion

5-azaC has long been recognized as a hypomethylating agent, with its clinical function largely attributed to transcriptional regulation^36^. Given that its RNA-dependent effects significantly altered the transcriptome, and that caRNA m^5^C depletion was more pronounced than tRNA m^5^C (Figure 1A), we next investigated whether and how the RNA-dependent process regulates transcription. Contrary to its long-term DNA hypomethylation effect, 5-azaC led to transcriptional downregulation across leukemia cell lines, as measured by 5-ethynyl uridine (EU) labeling of nascent RNA, after 24 hours of treatment (Figure 2A and S2A-E). Interestingly, despite the anticipated transcriptional upregulation due to DNA demethylation at 72 hours, RNA-dependent effects appeared to still dominate, and transcriptional levels remained decreased after 72 hours in all six cell lines tested. This aligns with previous reports of inconsistencies between differential gene expression and DNA methylation changes following 5-azaC treatment^9–12^, as well as clinical cases where some MDS patients benefited from 5-azaC treatment without significant DNA demethylation^20^. AML patient samples exhibited similar transcriptional downregulation after 24- and 72-hour 5-azaC treatment (Figure 2B and S2F-G), highlighting the clinical relevance of these RNA-dependent effects. Our results underscore the importance of the RNA-dependent mechanisms in 5-azaC-induced gene expression dysregulation.

**Figure 2.**
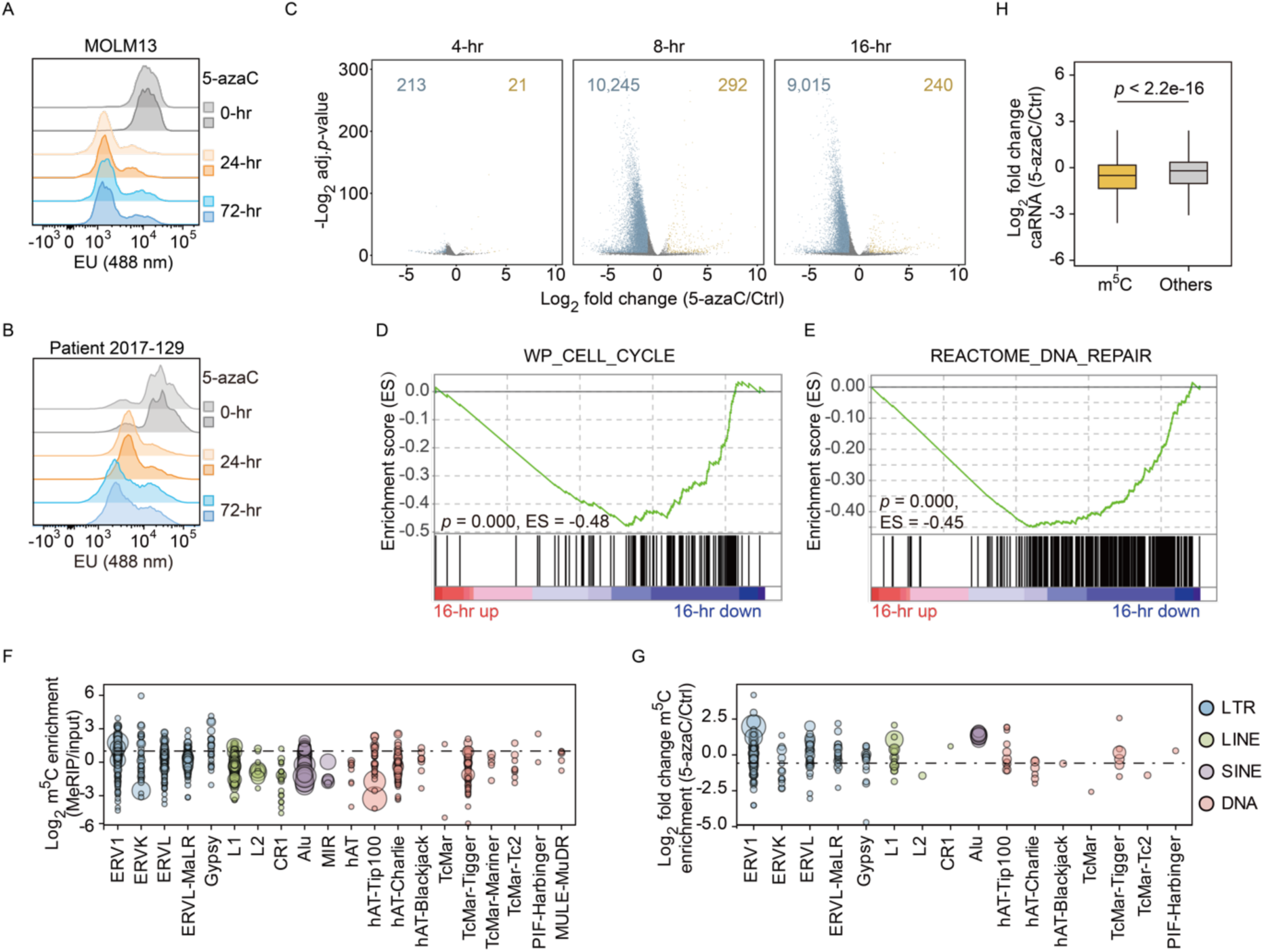
5-azaC induces transcriptional repression through caRNA m^5^C depletion. (A and B). EU fluorescent signals in MOLM13 cells (A) and AML patient sample 2017-129 (B) after 5-azaC (3 μM) treatment. (C) Nascent RNA-seq of MOLM13 cells after 3 μM of 5-azaC treatment. DEGs are defined as |log2 fold change (5-azaC/Ctrl) | > 1 and adj.*p*-value < 0.05. DEG numbers are annotated on the plot. Adj.*p*-value, adjusted *p*-value. (D and E) GSEA of transcriptional changes in cell cycle (D) and DNA repair (E) gene sets after 16 hours of 5-azaC (3 μM) treatment in MOLM13 cells. (F) m^5^C enrichment across repeat families in control MOLM13 cells. Circle sizes represent normalized read counts from MeRIP-seq. The dashed line indicates a value of 1. (G) Differential m^5^C enrichment across repeat families in MOLM13 cells after 16 hours of 5-azaC (3 μM) treatment. Circle sizes represent normalized MeRIP-seq read counts in untreated samples. The dashed line indicates a value of -0.58. (H) Differential levels of caRNA transcripts from genes adjacent to m^5^C sites (m^5^C) versus other genes (Others).

To assess gene-specific transcriptional changes, we performed nascent RNA-seq^30^ (Figure 2C and S2H), which captures newly synthesized RNA through biotin labeling of incorporated EU. The levels of nascent RNA were normalized to viable cell numbers by adding EU-labeled *Drosophila* spike-in. The nascent RNA-seq revealed significant transcriptional downregulation emerging from 8 hours of 5-azaC treatment, consistent with the starting time of the caRNA m^5^C decrease (Figure 1A) and a role of caRNA m^5^C in transcriptional activation^30^. GSEA confirmed transcriptional downregulation of cell cycle genes and DNA repair genes (Figure 2D-E and S2I-J), suggesting that the differential gene expression of corresponding pathways induced by 5-azaC is indeed derived from transcriptional regulation.

One of the DNA repair pathways, nucleotide excision repair (NER), is mediated by either global genome repair or transcription-coupled repair (TCR), with the latter dependent on transcriptional activity^37^. We reasoned that the overall transcriptional downregulation might contribute to the impaired DNA damage response caused by 5-azaC. Compared with controls, UDS after UVC irradiation was less susceptible to transcription inhibition in 5-azaC-treated cells (Figure S2K and L), implying a role of transcriptional suppression on the early DNA damage repair defects induced by 5-azaC.

To explore the mechanisms underlying transcriptional repression, we mapped m^5^C sites on caRNA using methylated RNA immunoprecipitation sequencing (MeRIP-seq) as previously reported^30^. RNA m^5^C levels were reduced after a 16-hour 5-azaC treatment in MOLM13 (Figure S2M), with 7,095 downregulated and 4,567 upregulated sites out of 21,158 peaks identified from control and treated samples. Notably, repetitive elements accounted for 62.6% of input reads and 88.27% MeRIP-enriched reads (Figure S2N), highlighting repeat RNA as a crucial group of caRNA that may exert transcriptional regulation. The increased proportion of repeat RNA in MeRIP samples further suggested preferential m^5^C installation. Similar to MOLM13, SKM1 cells also exhibited significant caRNA m^5^C downregulation after 16 hours of 5-azaC treatment (Figure S2O), with repeat RNA preferentially enriched in m^5^C MeRIP-seq datasets (Figure S2P). Further analysis identified long terminal repeat (LTR) elements as the most modified repeat RNA (Figure 2F), consistent with our previous findings^30^, while some subfamilies of long interspersed nuclear element (LINEs), short interspersed nuclear elements (SINEs), and DNA transposons were also m^5^C modified. 5-azaC treatment caused reduced m^5^C levels among most repeat RNAs (Figure 2G), consistent with the reduced overall caRNA m^5^C level measured with UHPLC-MS/MS (Figure 1A). RNA modifications on repeat RNA have emerged as key elements in the regulation of local chromatin state and transcription^30,38–40^, and caRNA m^5^C have been associated with maintaining an open chromatin state^30^. Therefore, the 5-azaC-induced m^5^C downregulation may drive the observed transcriptional repression. Indeed, 5-azaC treatment caused the downregulation of caRNA repeat subfamilies bearing m^5^C, particularly in the LTR family (Figure S2Q). Furthermore, genes adjacent to m^5^C sites exhibited greater downregulation than those farther away (Figure 2H), reinforcing the link between m^5^C loss and transcriptional repression of local chromatin regions.

### 5-azaC induces transcriptional repression through the m^5^C-dependent H2AK119ub regulation

We have recently shown that caRNA m^5^C regulates transcription by recruiting the m^5^C-binding protein MBD6 and its interacting partner, the PR-DUB deubiquitylase complex (BAP1/ASXL1/KDM1B complex), which mediates removal of the histone modification H2AK119ub to promote transcriptional activation^30^. Inhibition of RNA methyltransferase that installs caRNA m^5^C by 5-azaC would reduce PR-DUB recruitment and repress transcription. Indeed, we observed an increase in the global H2AK119ub level following a 16-hour 5-azaC treatment by immunoblotting (Figure 3A). H2AK119ub CUT&Tag analysis demonstrated elevated H2AK119ub levels near genomic regions of m^5^C-modified caRNA (Figure 3B-C). Consistent with the previously suggested role of LTRs in H2AK119ub regulation^30^, we observed the most significant upregulation of H2AK119ub at specific LTR subfamilies (Figure 3D). This upregulation was detected across gene loci and was particularly evident at transcription start sites (TSS) (Figure S3A), supporting its role in transcriptional regulation. Particularly, we observed enrichment of gene sets related to cell cycles and DNA damage repair pathways (Figure S3B), supporting that H2AK119ub changes contribute to the transcriptional repression of these pathways. Similar upregulation of H2AK119ub was also observed in 5-azaC-treated LK cells (Figure 3E). Cross-linking immunoprecipitation followed by high-throughput sequencing (CLIP-seq) revealed that 5-azaC induced decreased MBD6 binding at m^5^C sites (Figure 3B and S3C), consistent with the role of MBD6 as an m^5^C-binding protein involved in H2AK119ub regulation.

**Figure 3.**
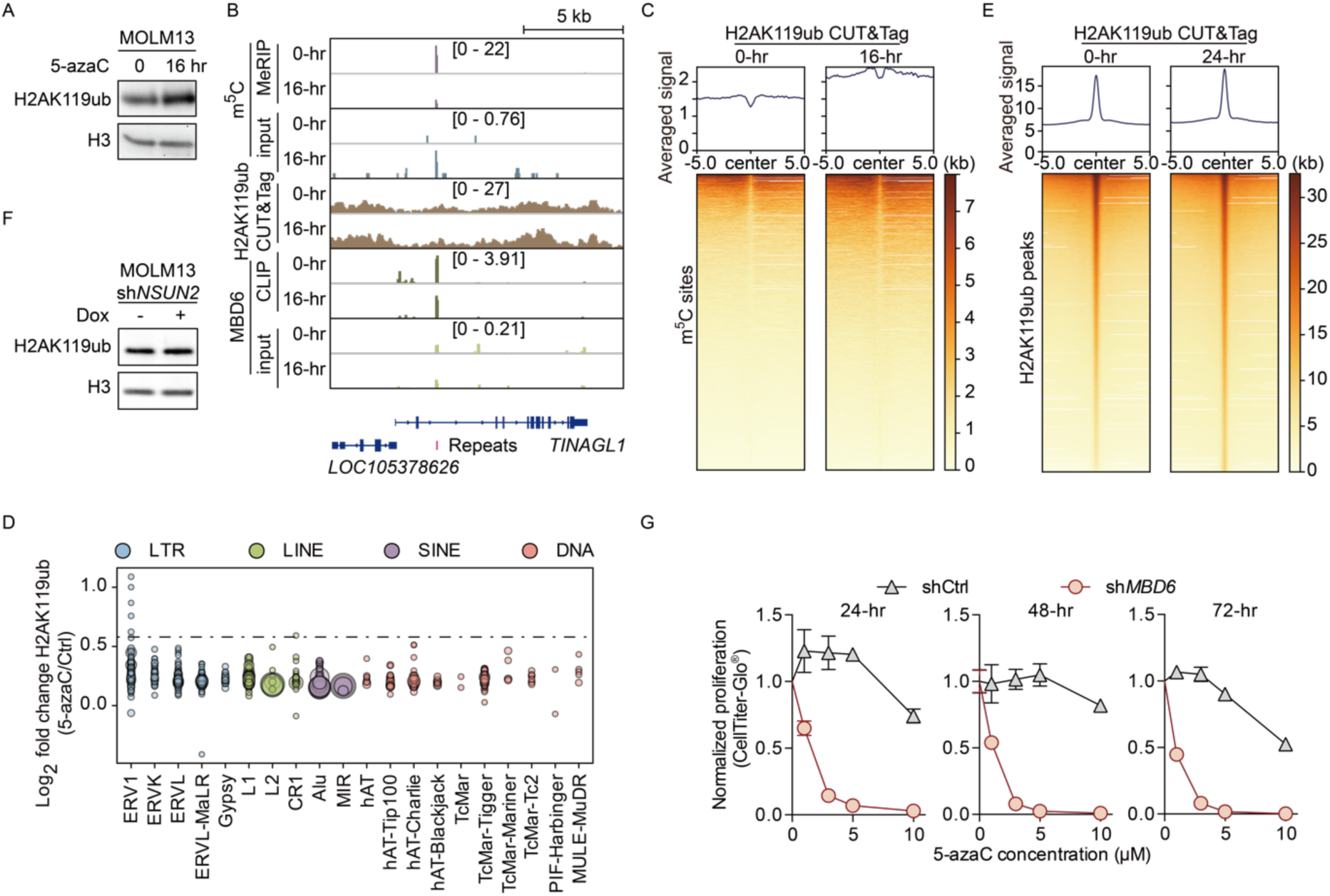
5-azaC induces transcriptional repression through m^5^C-dependent H2AK119ub regulation. (A) Immunoblotting of H2AK119ub changes after a 16-hour 5-azaC (3 μM) treatment in MOLM13 cells. (B) Integrative Genome Viewer (IGV) tracks showing m^5^C MeRIP-seq, H2AK119ub CUT&Tag, and MBD6 CLIP-seq signals at the *TINAGL1* locus after a 16-hour 5-azaC (3 μM) treatment. Track scales are indicated in brackets. (C) Metaplot and heatmap of H2AK119ub level changes at m^5^C sites after 5-azaC treatment (3 μM) in MOLM13 cells. (D) H2AK119ub changes across repeat families. Circle sizes represent normalized CUT&Tag read counts in control samples. The dashed line indicates a value of 0.58. (E) Metaplot and heatmap of H2AK119ub level changes at identified peaks after 3 μM of 5-azaC treatment in LK cells. (F) Immunoblotting of H2AK119ub changes following doxycycline (Dox)-induced NSUN2 depletion for 5 days in MOLM13 cells. (G) MOLM13 cell proliferation following 5-azaC treatment after 48-hour shRNA-mediated MBD6 depletion. Time points indicate hours after 5-azaC treatment.

To further validate the role of m^5^C in this transcriptional regulation, we depleted NSUN2, the primary methyltransferase responsible for installing m^5^C on caRNA^30^. *NSUN2* knockdown (KD) resulted in H2AK119ub upregulation (Figure 3F), supporting a causal role for m^5^C in this regulation. Notably, depletion of other m^5^C methyltransferases, including NSUN5, NOP2, NSUN6, NSUN7, and TRDMT1, did not induce similar H2AK119ub upregulation (Figure S3D and E), indicating that NSUN2 plays the primary role in the m^5^C-mediated regulation of H2AK119ub. Furthermore, depletion of the DNA 5mC methyltransferase DNMT1 led to a reduction in H2AK119ub levels, reinforcing that the early time point effect of 5-azaC on H2AK119ub goes through the RNA-dependent transcriptional repression rather than DNA hypomethylation.

MBD6 depletion has been shown to rescue the differentiation delay defect of *Tet2* deletion hematopoietic stem and progenitor cells (HSPCs) and inhibit leukemogenesis in *TET2* mutant leukemic cells, implying a potential role for MBD6 inhibition in leukemia treatment^30^. Consistently, *MBD6* KD sensitized MOLM13 cells to 5-azaC treatment (Figure 3G), indicating that combining 5-azaC with an MBD6 inhibitor could be a promising therapeutic approach for treating leukemia.

Overall, our findings support that 5-azaC represses transcription through caRNA m^5^C depletion that regulates transcription through the m^5^C/MBD6/PR-DUB/H2AK119ub axis.

### 5-azaC induces transcriptional repression through downregulation of H3K27ac

It has been reported that inhibitors of CBP/p300 exhibited a synergistic effect with 5-azaC but not with decitabine^41^, suggesting a link between the RNA-dependent function of 5-azaC and H3K27ac regulation. We indeed observed overall decreased H3K27ac levels in MOLM13 and SKM1 cells after a 16-hour 5-azaC treatment (Figure 4A and S4A). This observation suggested that H3K27ac downregulation may also contribute to the m^5^C-dependent transcriptional repression by 5-azaC. The reduction of H3K27ac persisted at later time points (Figure S4A and B), aligning with the predominant transcriptional repression effect of 5-azaC over DNA hypomethylation. H3K27ac CUT&Tag analysis confirmed a global downregulation of H3K27ac (Figure 4B and S4C), particularly affecting gene sets related to cell cycles and DNA damage repair (Figure 4C and S4D). To elucidate the relationship between H3K27ac downregulation and transcriptional repression, we grouped genes into four quantiles based on the degree of the adjacent H3K27ac peak changes, with Q1 representing the most downregulated genes and Q4 the most upregulated.

**Figure 4.**
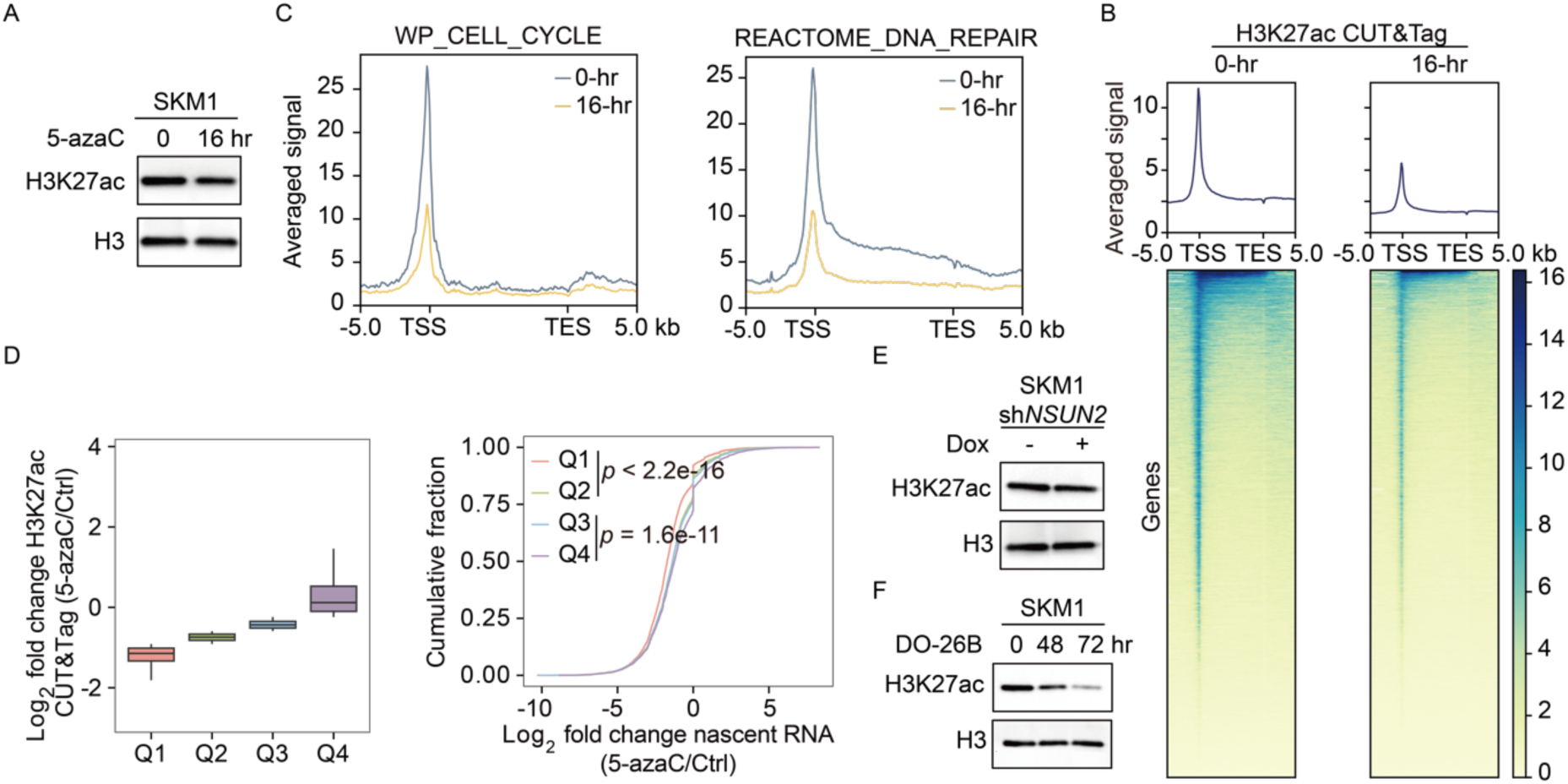
5-azaC induces transcriptional repression through H3K27ac downregulation. (A) Immunoblotting of H3K27ac changes in SKM1 cells after a 16-hour 5-azaC (3 μM) treatment. (B) Metaplot and heatmap of H3K27ac changes across all annotated genes after a 16-hour 5-azaC (3 μM) treatment in SKM1 cells. (C) Metaplot of H3K27ac changes in cell cycle and DNA damage repair gene sets in SKM1 cells. (D) H3K27ac (left) and nascent RNA-seq (right) changes in genes grouped into four quantiles based on fold changes of nearby H3K27ac peaks in SKM1 cells. Q1 represents the quantile with the most negative fold change. (E and F) Immunoblotting of H3K27ac changes after a 5-day Dox-induced depletion of NSUN2 (E) or 5 μM DO-26B treatment (F) in SKM1 cells.

A comparison with nascent RNA-seq data revealed a correlation between H3K27ac loss and transcriptional repression in downregulated genes (Figure 4D and S4E). Most H3K27ac downregulation occurred in regions enriched with LTRs and LINEs (Figure S4F and G). Decreased H3K27ac was observed at genomic loci corresponding to m^5^C-modified sites in both MOLM13 and SKM1 (Figure S4H and I), supporting RNA m^5^C-dependent H3K27ac downregulation by 5-azaC treatment.

To further elucidate the causal relationship between m^5^C and H3K27ac, we examined NSUN2, the primary caRNA m^5^C methyltransferase^30^. NSUN2 depletion caused a reduction of the H3K27ac level (Figure 4E and S4J). Among other m^5^C methyltransferases, we observed a subtle decrease of H3K27ac following *NSUN7* KD (Figure S4J), consistent with its reported role in eRNA m^5^C modification^42^. Depleting NSUN5, NOP2, NSUN6, or TRDMT1 did not induce H3K27ac reduction, suggesting their RNA targets are not involved in this regulation. Notably, depletion of DNMT1 resulted in H3K27ac upregulation, further confirming that the early time point H3K27ac disruption observed with 5-azaC treatment is RNA-dependent rather than DNA-dependent. The catalytic activity of NSUN2 is responsible for the regulation, as revealed by H3K27ac downregulation of cells treated with NSUN2 inhibitor DO-26B^43^ (Figure 4F). We performed H3K27ac CUT&Tag to explore genomic effects of NSUN2 depletion, which revealed H3K27ac downregulation at m^5^C sites (Figure S4K), further supporting the causal relationship between m^5^C and H3K27ac.

### 5-azaC downregulates H3K27ac through suppressing SRSF2-mediated p300 recruitment

To elucidate the underlying mechanism connecting caRNA m^5^C to H3K27ac, we examined a nuclear localized m^5^C-binding protein, SRSF2^26^. SRSF2 depletion resulted in a significant reduction of H3K27ac (Figure 5A and S5A-B), suggesting its role in H3K27ac regulation. Given that SKM1 cells exhibited significantly higher SRSF2 expression than MOLM13 cells (Figure S5C), we selected SKM1 to further explore SRSF2-dependent H3K27ac regulation. Previous studies have shown that SRSF2 associates with p300, thereby influencing H3K27ac deposition^44,45^. Consistently, immunofluorescence analysis revealed colocalization of SRSF2 and p300 (Figure 5B and C). Proximity ligation assay (PLA) confirmed their interaction (Figure 5D), supporting that SRSF2 recruits p300 to regulate H3K27ac deposition.

**Figure 5.**
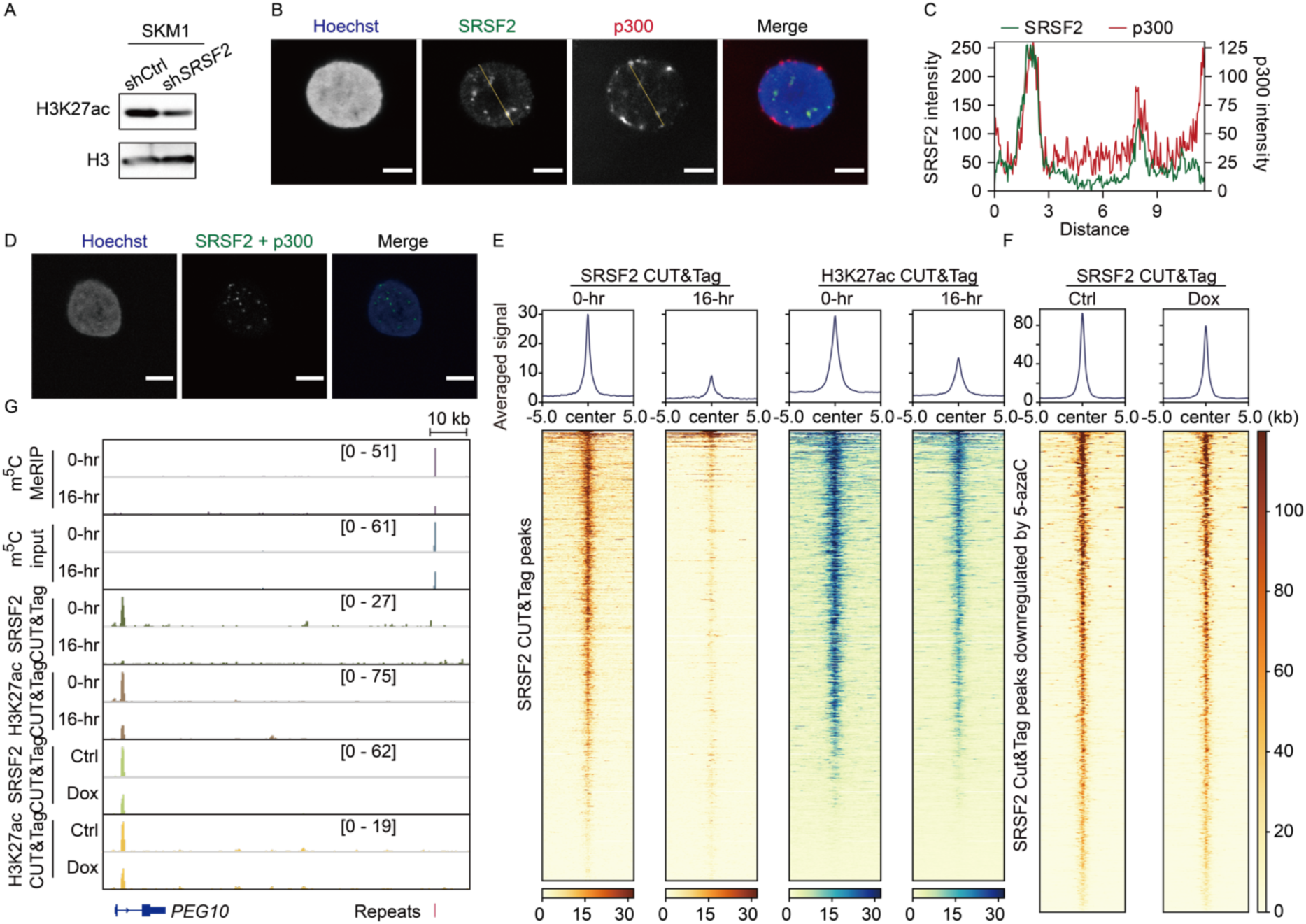
5-azaC downregulates H3K27ac through the SRSF2-mediated p300 recruitment. (A) Immunoblotting of H3K27ac changes after a 48-hour shRNA-mediated SRSF2 depletion in SKM1 cells. (B) Co-immunofluorescence of SRSF2 and p300 in SKM1 cells. Scale bar: 5 μm. (C) Quantification of SRSF2-p300 colocalization along the yellow line in (B). (D) PLA of SRSF2 and p300 in SKM1 cells. Scale bar: 5 μm. (E) Metaplot and heatmap of SRSF2 and H3K27ac changes at SRSF2 target sites after 5-azaC (3 μM) treatment. (F) Metaplot and heatmap of SRSF2 CUT&Tag level changes at downregulated SRSF2 peaks after 5-azaC (3 μM) treatment. Samples were treated with Dox for 5-day NSUN2 depletion. (G) IGV tracks showing m^5^C MeRIP-seq, SRSF2 CUT&Tag, and H3K27ac CUT&Tag signals at genomic locus near *PEG10* after a 16-hour 5-azaC (3 μM) treatment or Dox-induced NSUN2 depletion. Track scales are indicated in brackets.

Based on these observations, we hypothesized that 5-azaC disrupts m^5^C-guided genomic binding of SRSF2, thereby impairing p300 localization and H3K27ac installation. Due to the inability of commercial SRSF2 antibodies to efficiently immunoprecipitate SRSF2 (Figure S5D), we overexpressed SRSF2-Myc in K562 cells to assess its RNA binding following m^5^C-depletion by 5-azaC. After 16 hours of treatment, SRSF2 exhibited reduced RNA binding, as detected by cross-linking immunoprecipitation (CLIP) with end ligation of pCp-AF488 fluorophore (Figure S5E). Given that SRSF2 binding to chromatin is known to be RNA-dependent^46^, we speculated that the impaired RNA binding of SRSF2 leads to reduced chromatin binding of SRSF2 and subsequent loss of H3K27ac deposition. To test this, we performed SRSF2 CUT&Tag after 16 hours of 5-azaC treatment. We indeed observed a significant reduction in SRSF2 chromatin binding, which correlated with decreased H3K27ac (Figure 5E and G). SRSF2 was highly enriched at promoters of genes involved in cell cycle and DNA damage repair pathways, and its binding at these sites was diminished following 5-azaC treatment (Figure S5F). This is consistent with the recent findings that genes involved in DNA damage repair are particularly sensitive to transcriptional regulation by SRSF2^47^. Furthermore, SRSF2 binding was downregulated at chromatin regions adjacent to m^5^C sites (Figure S5G), supporting the model in which caRNA m^5^C facilitates SRSF2 recruitment and H3K27ac deposition. Depletion of caRNA m^5^C would therefore reduce nearby H3K27ac.

To establish a direct link between m^5^C and SRSF2 chromatin binding, we performed SRSF2 CUT&Tag after NSUN2 depletion in SKM1 cells. SRSF2 chromatin binding was reduced at regions where its occupancy was also downregulated by 5-azaC (Figure 5F and G), reinforcing the role of m^5^C in guiding SRSF2 localization to chromatin. The reduction was most pronounced at TSS (Figure S5H), particularly for genes involved in cell cycle and DNA damage repair pathways (Figure S5I), aligning with early transcriptional changes induced by 5-azaC.

In summary, in addition to the m^5^C/MBD6/PR-DUB/H2AK119ub pathway, we uncovered a parallel pathway of 5-azaC-induced transcriptional repression through the m^5^C/SRSF2/p300/H3K27ac axis, highlighting RNA m^5^C as the critical link in epigenetic regulation and transcriptional repression effect of 5-azaC.

### NSUN2 regulates leukemia cell proliferation and hematopoietic lineage commitment

Given that NSUN2 is the primary m^5^C methyltransferase target responsible for the 5-azaC-mediated transcriptional repression, NSUN2 depletion should recapitulate the reduced proliferation of leukemic cells observed with 5-azaC treatment. Indeed, we observed proliferation inhibition after NSUN2 depletion in MOLM13 cells (Figure 6A). This aligns with previous reports in AML and other cancer cell types, where NSUN2 depletion inhibits cell proliferation^48–50^. GSEA of RNA-seq datasets from *NSUN2* KD MOLM13 cells revealed a downregulation of cell cycle genes (Figure 6B). NSUN2 depletion led to a decreased S-phase population in MOLM13 and SKM1 cells (Figure 6C and S6A), indicating prolonged cell cycle progression. Additionally, GSEA also indicated a downregulation of DNA damage repair pathway genes (Figure 6D). We validated the reduced expression of key DNA damage repair regulators CHK1 and RAD51 through immunoblotting (Figure 6E). Upon UVC irradiation, NSUN2 depletion caused diminished γH2A.X signaling (Figure 6F) and defective DNA repair activity, as demonstrated by EdU incorporation followed by click chemistry (Figure 6G and S6B). The cytotoxicity of 5-azaC could be partially rescued by NSUN2 overexpression (Figure S6C), reinforcing the role of NSUN2 in the RNA-dependent effect of 5-azaC.

**Figure 6.**
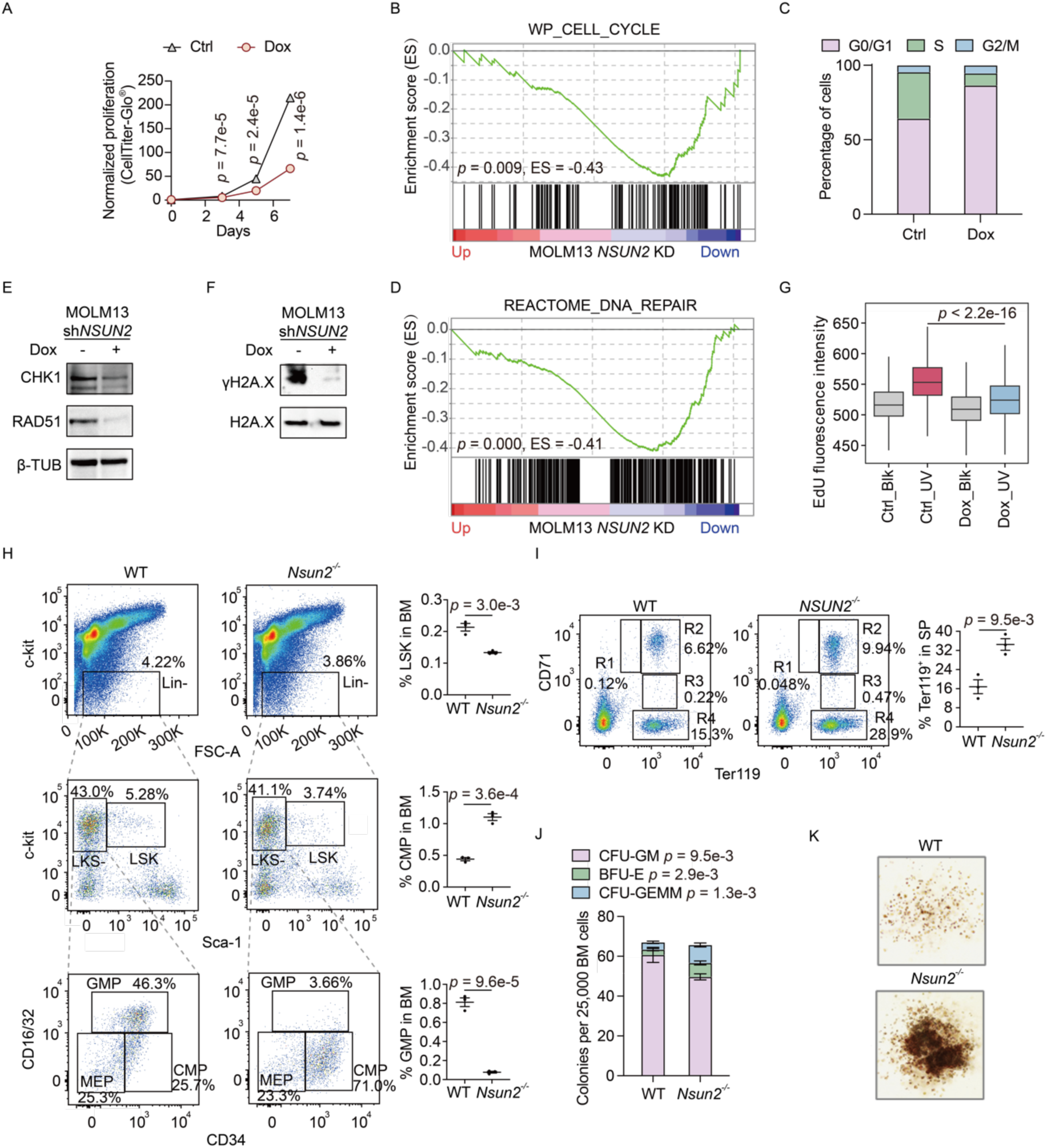
NSUN2 regulates leukemia proliferation and hematopoietic lineage commitment. (A) Cell proliferation of MOLM13 cells after Dox-induced NSUN2 depletion. (B) GSEA of MSigDB cell cycle gene set in MOLM13 cells after a 5-day Dox-induced NSUN2 depletion. (C) Cell cycle distribution of MOLM13 cells after a 5-day Dox-induced NSUN2 depletion. (D) GSEA of MSigDB DNA repair gene set in MOLM13 cells after a 5-day Dox-induced NSUN2 depletion. (E) Immunoblotting of DNA repair genes in MOLM13 cells after a 5-day Dox-induced NSUN2 depletion. (F) Immunoblotting of γH2A.X in MOLM13 cells after a 5-day Dox-induced NSUN2 depletion followed by 50J/m^2^ UVC irradiation. (G) EdU incorporation during DNA damage repair after UVC irradiation in MOLM13 cells after a 5-day Dox-induced NSUN2 depletion. Only G0/G1-phase populations are shown. Blk, no irradiation. (H) Flow cytometry analysis of LSK, LKS^-^, GMP, MEP, and CMP populations in BM from 6-month-old wild-type (WT) and *Nsun2*^-/-^ mice (left), and quantification of LSK, CMP and GMP populations in the total BM cells (right). N = 3 mice/genotype. (I) Flow cytometry analysis of erythroid cells in the spleen of 6-month-old WT and *Nsun2*^-/-^ mice (left), and quantification of the Ter119^+^ erythroblasts (right). R1: pro-, R2: basophilic-, R3: late basophilic- and polychromatophilic-, and R4: orthochromatophilic-erythroblasts. SP, spleen. (J) CFU-C assay of BM cells from 6-month-old WT and *Nsun2*^-/-^ mice. (K) Representative GEMM colonies formed by BM cells from 6-month-old WT and *Nsun2*^-/-^ mice.

If NSUN2 is one of the key functional targets of 5-azaC, its depletion may mimic the RNA-dependent effects of 5-azaC during hematopoiesis. MDS are typically characterized by ineffective erythropoiesis^51^, and monocytosis has been associated with earlier transformation to AML^52^. The use of 5-azaC helps restore erythroid differentiation^53,54^. We investigated hematopoietic lineage commitment in *Nsun2* knockout (KO) mice. BM cells purified from 6-month-old *Nsun2* KO mice (Figure S6D) exhibited a reduced frequency of Lin^-^Sca1^+^c-Kit^+^ (LSK) cells (Figure 6H), indicative of impaired HSPC maintenance. We also observed a decrease in granulocyte-monocyte progenitor (GMP) cells and an increase in common myeloid progenitor (CMP) cells in *Nsun2* KO BM cells compared to wild type (WT) (Figure 6H). Additionally, *Nsun2* KO mice exhibited a higher population of Ter119^+^ erythroid cells in the spleen (Figure 6I), suggesting a shift toward erythropoiesis as reported when using 5-azaC in patients^53,54^. *In vitro* colony-forming unit cell (CFU-C) assays using *Nsun2* KO BM cells revealed a lower frequency and number of CFU-granulocyte/macrophage (CFU-GM) colonies, accompanied by an increase in burst-forming unit-erythroid (BFU-E) and CFU-granulocyte-erythrocyte-monocyte-megakaryocyte (CFU-GEMM) colonies (Figure 6J). This is consistent with the observed reduction in GMPs and expansion of CMPs in *Nsun2* KO mice. Furthermore, CFU-GEMM colonies derived from *Nsun2* KO BM cells were larger in size (Figure 6K), suggesting altered differentiation dynamics. The progeny of *Nsun2* KO CFU-C cultures exhibited a reduced Lin^-^c-Kit^+^ (LK) population (Figure S6E) and an increased proportion of Ter119^+^ erythroid cells (Figure S6F), further supporting a shift toward erythropoiesis with impeded myelopoiesis after *Nsun2* deletion.

Overall, our results demonstrate that NSUN2 depletion significantly promotes erythroid lineage commitment while suppressing granulocytic/monocytic differentiation, effects consistent with those observed with 5-azaC treatment.

### Correlation of leukemia cell line mutations with 5-azaC sensitivity

Leukemia cell lines exhibit varying sensitivity to 5-azaC (Figure 1B-E). While the DNA-dependent cytotoxicity of 5-azaC has been extensively studied^55–59^, the factors influencing sensitivity to its RNA-dependent effect remain poorly understood. To address this and draw a direct connection between the new pathways we uncovered with 5-azaC sensitivity, we screened 26 different leukemia cell lines for their response to 5-azaC at the early time point (Figure 7A). Consistent with our previous findings, a 24-hour decitabine treatment had little impact on proliferation, whereas 5-azaC led to varying levels of growth inhibition across different cell lines. We integrated publicly available mutation data and identified mutations enriched in 5-azaC-sensitive cells that are frequently observed in clinical MDS and AML samples. This includes *TET2*, which catalyzes the oxidation of RNA m^5^C^30^. This aligns with previous reports that MDS and AML patients harboring *TET2* mutations tend to respond more effectively to 5-azaC treatment^54,55,60^, and agrees with the m^5^C/MBD6/PR-DUB/H2AK119ub regulatory axis we reported recently^30^ and in this study. Consistently, *TET2* KO THP1 cells exhibited increased sensitivity to 5-azaC, particularly at early time points (Figure 7B), indicating that *TET2* mutation is likely a reliable biomarker for the 5-azaC sensitivity. Beyond *TET2*, mutations in *ASXL1*, a component of the PR-DUB complex, were also enriched in 5-azaC-sensitive cells.

**Figure 7.**
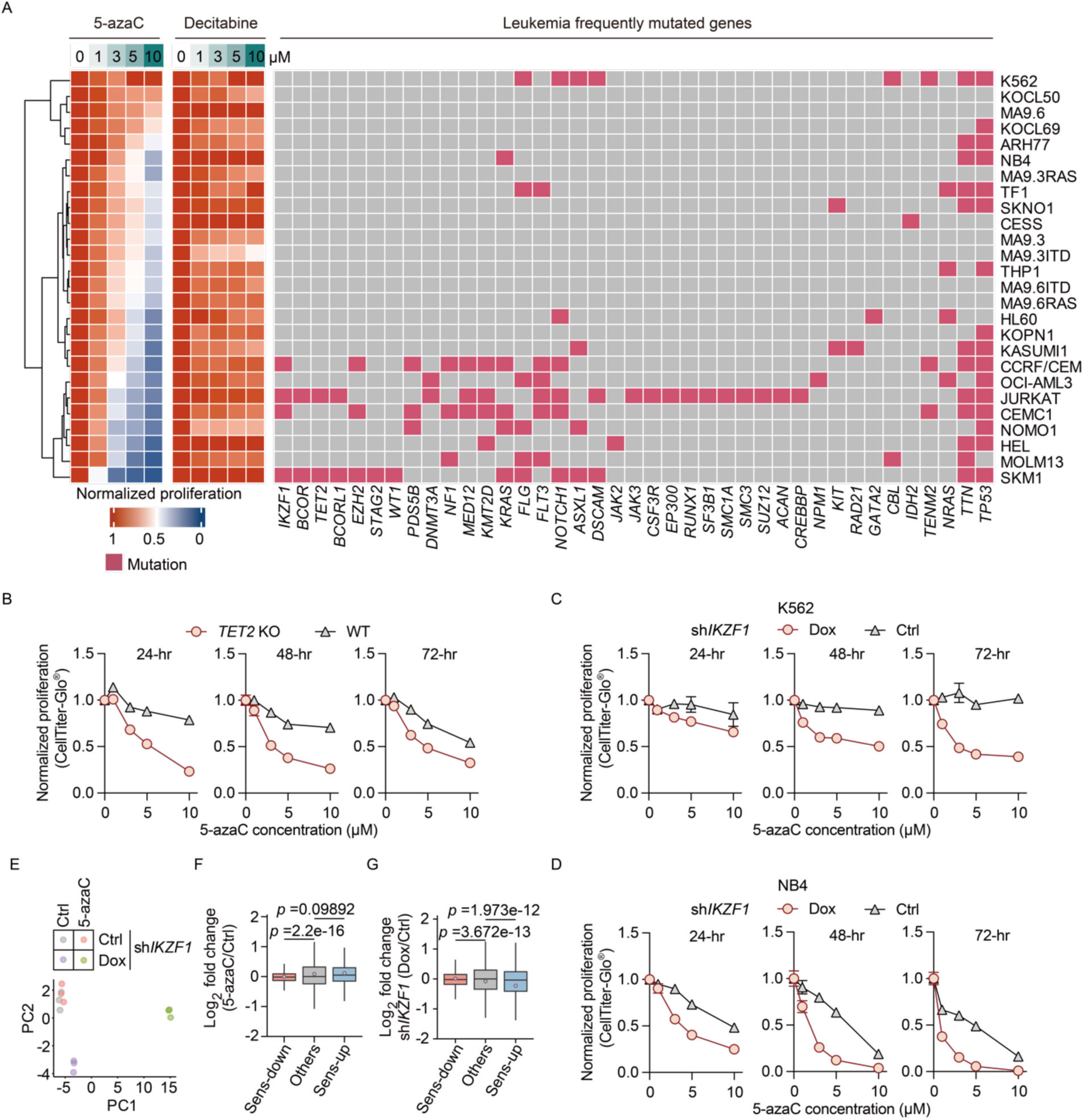
Screening of leukemia cell lines infers mutations associated with 5-azaC sensitivity. (A) Proliferation of 26 leukemia cell lines after a 24-hour treatment with 5-azaC (left) or decitabine (middle), along with their reported mutations from public databases (right). Proliferation was measured by CellTiter-Glo^®^. (B) Proliferation of THP1 WT and *TET2* KO cells after 5-azaC treatment. (C and D) Proliferation of K562 (C) and NB4 (D) cells treated with 5-azaC after Dox-induced IKZF1 depletion. The denoted time indicates hours after 5-azaC treatment. (E) PCA of RNA-seq of K562 cells with Dox-induced IKZF1 depletion and/or a 24-hour 5-azaC (3 μM) treatment (F and G) Differential expression of Sens-up, Sens-down and other genes (Others) in K562 cells with 5-azaC (3 μM) treatment without IKZF1 depletion (F) or Dox-induced IKZF1 depletion without 5-azaC treatment (G). The lilac circle represents value of mean in each group of genes.

In addition to RNA m^5^C-related pathways, we identified *IKZF1* as another potential class of functionally related mutations. While *IKZF1* mutations are more commonly observed in acute lymphoblastic leukemia (ALL)^61^, they have also been identified in AML as an independent marker of adverse risk with unique clinical phenotypes^62^. Moreover, IKZF1 degraders have shown efficacy in eliminating *MLL1*-rearranged (*MLL*-r) AML cells^63^. In our proliferation assay, IKZF1 depletion sensitized K562 cells (Figure 7C and S7A) and NB4 cells (Figure 7D) to 5-azaC treatment. RNA-seq revealed that treatment with 3 μM 5-azaC induced only subtle transcriptomic changes in K562 cells at 24 hours, consistent with its resistance to 5-azaC. In contrast, IKZF1 depletion markedly amplified the transcriptional response (Figure 7E). Genes significantly up- or down-regulated by 5-azaC in IKZF1-depleted K562 cells (denoted as Sens-up and Sens-down, respectively) showed only mild up- and down-regulated in control cells (Figure 7F). However, the Sens-up and Sens-down genes were notably down- and up-regulated after IKZF1 depletion (Figure 7G), suggesting that IKZF1 depletion may reprogram the transcriptomic landscape, rendering cells more reliant on Sens-down genes and more vulnerable to Sens-up gene activation, thereby increasing sensitivity to 5-azaC. IKZF1 ChIP-seq from the ENCODE database^64,65^ revealed enrichment of IKZF1 binding at genomic loci containing m^5^C sites previously identified in K562 cells^30^ (Figure S7B), suggesting a functional interplay between IKZF1 and RNA m^5^C-mediated regulation.

To further investigate potential drivers of 5-azaC sensitivity, we applied the network inference algorithm NetBID2^66^ to model regulatory activities of frequently mutated genes in MDS and AML. We constructed a leukemia-specific gene regulatory network using SJARACNe algorithm, employing RNA-seq datasets of 72 myeloid-origin leukemia cell lines from the DepMap portal^67^. After overlaying this network with the five leukemia cell lines most sensitive to 5-azaC (SKM1, MOLM13, HEL, NOMO1, OCI-AML3) and six most 5-azaC resistant ones (K562, NB4, TF1, SKNO1, THP1, HL60), we assessed hub gene activity based on two criteria: 1) differential expression between sensitive and resistant cell lines; and 2) differential expression of target genes between the two groups (Figure S7C). This analysis identified three candidate drivers of 5-azaC sensitivity: *SRSF2*, *NPM1*, and *EZH2* (Figure S7D-E). The expression levels of these genes and their targets in leukemic cells correlate with 5-azaC sensitivity (Figure S7D). Positively regulated target genes of these candidate drivers are expressed at lower levels in sensitive cell lines, while negatively regulated targets are more highly expressed (Figure S7E). Additionally, *EZH2* mutations were enriched in 5-azaC-sensitive leukemic cells (Figure 7A). Functional validation revealed that *NPM1* KD increased THP1 sensitivity to 5-azaC (Figure S7F), and pretreatment of HEL cells with EZH2 inhibitor tazemetostat led to a modest increase in 5-azaC sensitivity (Figure S7G).

The identification of SRSF2 as a potential driver to 5-azaC sensitivity from our network analysis further supports the RNA m^5^C-dependent H3K27ac deposition pathway mediated through the SRSF2/p300 axis we uncovered in this study. Additionally, NSUN2 inhibition would reduce caRNA m^5^C level, leading to increased H2AK119ub^30^, which is known to recruit PRC2 for H3K27me3 deposition^68^. This could synergize with reduced chromatin binding of SRSF2 and H3K27ac downregulation. The *EZH2* mutations may also be associated with this RNA-dependent pathway, which requires future studies. Overall, our leukemia cell line screen and network analysis identified new candidate mutations that may serve as biomarkers for predicting 5-azaC sensitivity or as potential therapeutic targets for combination treatments.

## DISCUSSION

As a ribonucleoside analogue, 5-azacytidine (5-azaC) can be incorporated into both nascent RNA and nascent DNA to covalently inhibit RNA and DNA methyltransferases. While traditionally regarded as a DNA hypomethylating agent with consolidated DNA hypomethylation effects after a period of time, 5-azaC leads to differential gene expression that could not be explained solely by its DNA hypomethylation changes^9–12^. Notably, 5-azaC must be converted into decitabine to be incorporated into DNA, whereas more than 80% of 5-azaC is directly incorporated into RNA, inducing more significant inhibition of RNA m^5^C installation. Here we demonstrate that: i) 5-azaC treatment causes RNA m^5^C hypomethylation within the first 24 hours, and this effect is sufficient to drive cytotoxicity in leukemic cells, independent of DNA hypomethylation; ii) the RNA m^5^C hypomethylation explains the gene repression effect of 5-azaC; iii) we identify two pathways responsible for the observed gene repression by 5-azaC. Depletion of m^5^C in caRNA disrupts the MBD6-mediated H2AK119ub deubiquitination, leading to globally elevated H2AK119ub levels. In parallel, it impairs SRSF2 binding to chromatin, reducing p300 recruitment for H3K27ac deposition; iv) NSUN2 is the primary target responsible for 5-azaC-induced gene repression, and our animal model studies demonstrate that Nsun2 depletion in HSPCs causes hematopoietic lineage commitment towards erythroid, reminiscent of that also observed in patients treated with 5-azaC^53,54^ ; and lastly v) our leukemia cell line screen identified *TET2* or *IKZF1* mutations that can sensitize 5-azaC treatment, consistent with the RNA-dependent cytotoxicity of 5-azaC in leukemic cells. In summary, our findings highlight the transcription repression effect of 5-azaC through caRNA m^5^C depletion, providing insights into its mechanism of action, predictors of treatment efficacy, and future directions for therapy developments based on 5-azaC.

Some patients benefited from 5-azaC treatment did not exhibit DNA demethylation^20,69^, and hypomethylated samples displayed differential gene expression patterns that could not be explained by DNA methylation changes^9–12^. By examining transcriptional dysregulation at early time points, we demonstrated that the NSUN2-mediated RNA methylation pathways are responsible for transcriptional repression observed with 5-azaC treatment. The RNA-dependent effect may be more transient due to the dynamic nature of RNA synthesis and degradation, in contrast to the heritable stability of DNA methylation. This potentially explains the necessity for continuous administration of 5-azaC in treatment^70^. The opposing effects of RNA- and DNA-dependent transcriptional regulation can be complicated and may amplify or attenuate each other depending on the genetic backgrounds of recipient individuals. Patient cohorts may selectively respond to either RNA methylation or DNA methylation inhibition. It could be beneficial to develop small molecule inhibitors that selectively target the RNA m^5^C-dependent pathway for more precise therapies.

Mechanistically, our studies revealed that the RNA-dependent transcriptional repression by 5-azaC leads to the downregulation of cell cycle and DNA damage repair genes, impairing leukemia cell survival, increasing cellular stress, and promoting apoptosis. This also supports the contribution of RNA-dependent transcription suppression pathway to the enhanced efficacy of 5-azaC when combined with venetoclax. The transcriptional repression by 5-azaC can be mediated by at least two pathways: the m^5^C/MBD6/PR-DUB/H2AK119ub axis and the m^5^C/SRSF2/p300/H3K27ac axis. The elevated H2AK119ub and reduced H3K27ac synergize with each other to repress transcription. Notably, we observed persistent H3K27ac reduction during prolonged 5-azaC treatment. H3K27ac is associated with actively transcribed genes, and its local chromatin environment may be less impacted by the DNA hypomethylating effects of 5-azaC in longer time points, leading to a sustained decrease in transcriptional activity. The varying contributions of H2AK119ub and H3K27ac regulation across heterogeneous leukemia cases may serve as predictive markers for the relative dominance of DNA- and RNA-dependent effects during long-term treatment.

This study primarily focused on the caRNA-dependent transcriptional repression by 5-azaC, but its effects on other RNA species (mRNA, rRNA, and tRNA) should be explored in the future. We show that NSUN2 is the primary enzyme responsible for the caRNA demethylation and transcription repression caused by 5- azaC treatment. We have carefully examined *Nsun2* knockout mice and observed shifted hematopoietic lineage commitment towards erythropoiesis, consistent with the role of 5-azaC in restoring erythroid differentiation^53,54^.

The heterogeneity of myeloid malignancies caused varying responses to 5-azaC treatment. A previous study comparing 5-azaC-sensitive and -resistant cell lines found differences in RNA m^5^C methyltransferase levels and their related transcriptional regulation^21^. Our study fills a critical missing piece in understanding the RNA-dependent transcription regulation by 5-azaC, which leads to cell cycle arrest and attenuated DNA damage repair. Consistent with the importance of the RNA-dependent function, our leukemia cell line screen and gene network inference revealed mutations related to RNA m^5^C regulation, including *TET2*, *ASXL1*, and *SRSF2*. These findings illustrated the critical role of the RNA-dependent mechanisms in determining 5-azaC sensitivity. Additionally, we identified EZH2, the methyltransferase for H3K27me3, which is closely linked to the regulation of H2AK119ub^71^ described above. We further confirmed *TET2* as a predictive biomarker for 5-azaC sensitivity. In parallel, we revealed that IKZF1 depletion sensitizes leukemic cells to 5-azaC, suggesting IKZF1 as a potential new target to synergize with 5-azaC.

In summary, our study unveils that 5-azaC mediates transcriptional repression through RNA m^5^C-dependent regulation of H2AK119ub and H3K27ac. The RNA-dependent pathways disrupt cell cycle and DNA damage repair pathways, contributing to 5-azaC-induced cytotoxicity, and play crucial roles in modulating the sensitivity of myeloid malignant cells to 5-azaC.

## RESOURCE AVAILABILITY

### Lead contact

Further information and requests for resources and reagents should be directed to and will be fulfilled by the lead contact, Chuan He.

### Materials availability

All the materials generated in this manuscript are available from the lead contact under a complete Materials Transfer Agreement.

### Data and code availability

- Sequencing data generated from this study have been deposited in the Gene Expression Omnibus (GEO) and made accessible under accession numbers GEO: GSE297589 and GSE297591
- This paper does not report original code.
- Any additional information required to reanalyze the data reported in this paper is available from the lead contact upon request.

## ACKNOWLEDGMENTS

We thank Dr. J. Chen at Beckman Research Institute for providing the leukemia cell lines for 5-azaC sensitivity screening. We thank Dr. L Wang at Northwestern University for providing the MBD6 antibodies for CLIP-seq. Funding: This work is supported by the National Institute of Health R01 HL174477 (C.H. and M.X.), the Cancer Prevention & Research Institute of Texas (CPRIT RP250055 to M.X.) and the Ludwig Center for Metastasis at the University of Chicago. C. H. is an investigator of the Howard Hughes Medical Institute.

## AUTHOR CONTRIBUTIONS

Conceptualization, C.H., B.G., and M.X.; methodology, Z.Z.; Investigation, B.G., Y.L., L.Z., Z.Z., X.M., J.X., F-C.Y. and X.D.; writing—original draft, C.H. and B.G.; writing—review & editing, C.H., M.X., F-C.Y., B.G. and Y.L.; funding acquisition, C.H. and M.X.; resources, M.X., J.H. and Y.L.; supervision, C.H. and M.X.

## SUPPLEMENTAL INFORMATION

**Document S1. Figures S1–S7**

**Table S1. Sequences of qPCR primers and shRNA**

**Table S2. Differential expression of genes after 24-hour 5-azaC treatment, related to Figure 1**

**Table S3. Genetic background of the AML patients in this study, related to Figure 2**

**Table S4. Differential transcriptional rate of genes after early 5-azaC treatment, related to Figure 2**

**Table S5. Differential expression of genes after *NSUN2* depletion in MOLM13, related to Figure 6**

## STAR★METHODS

### KEY RESOURCES TABLE

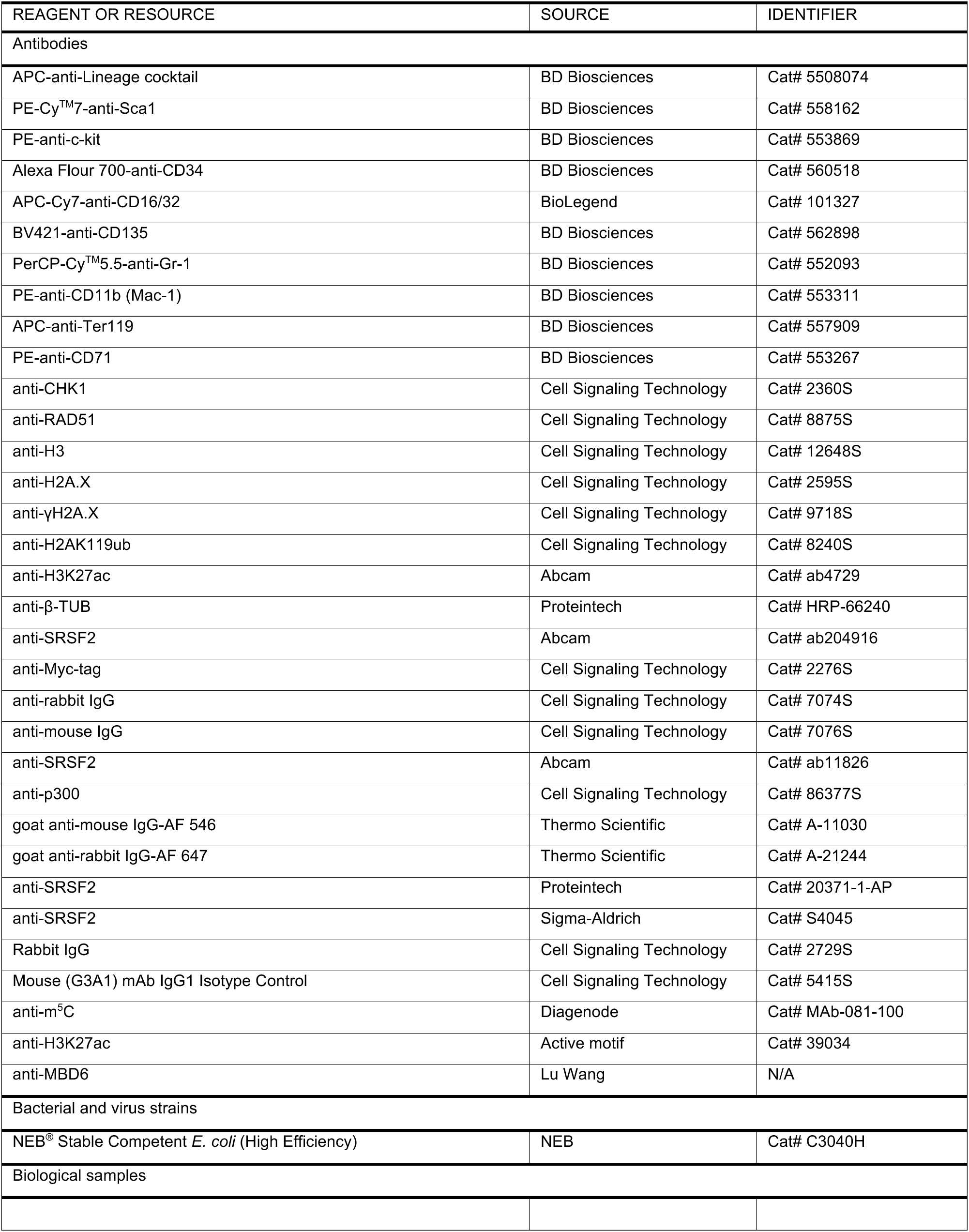
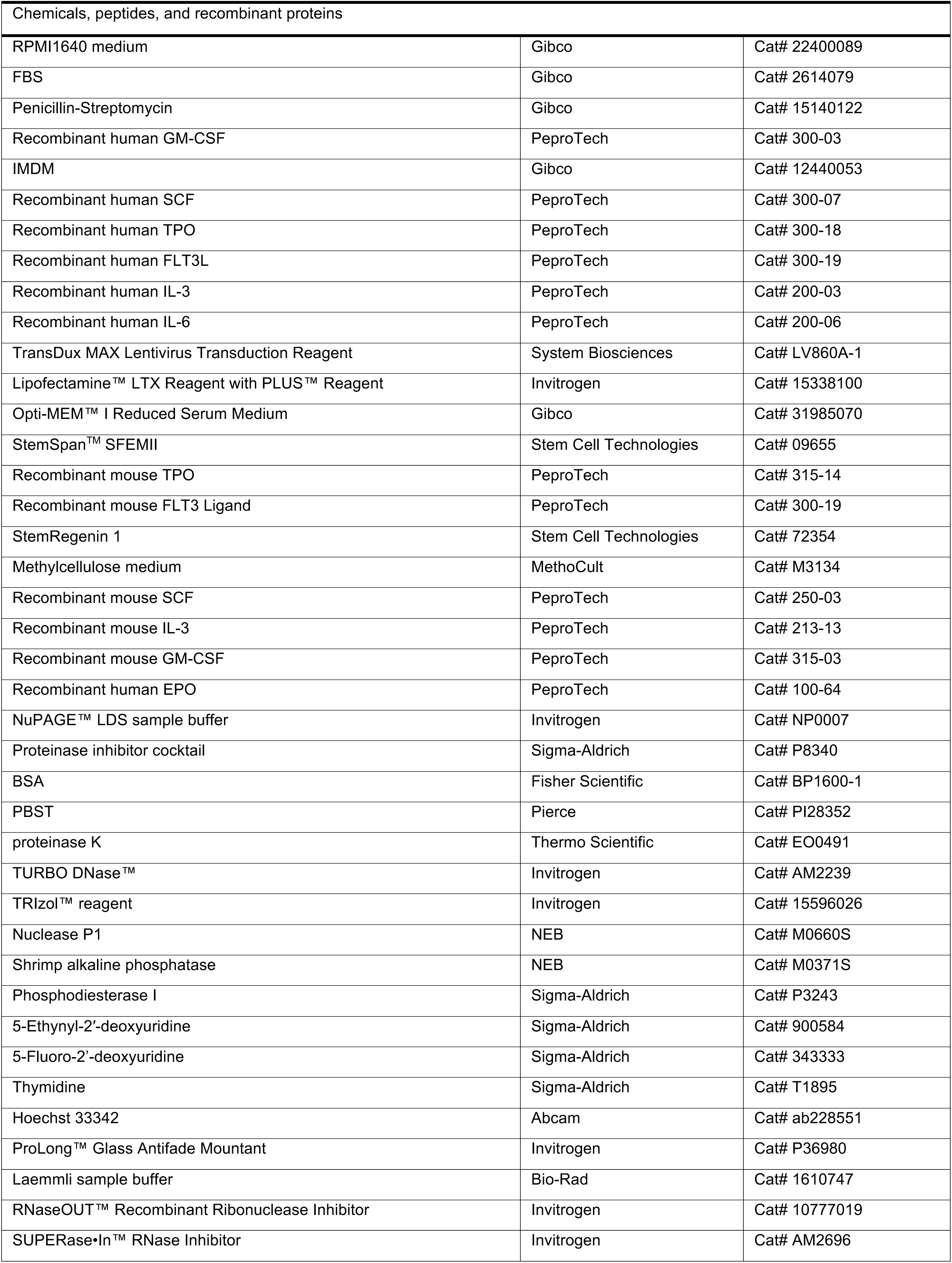
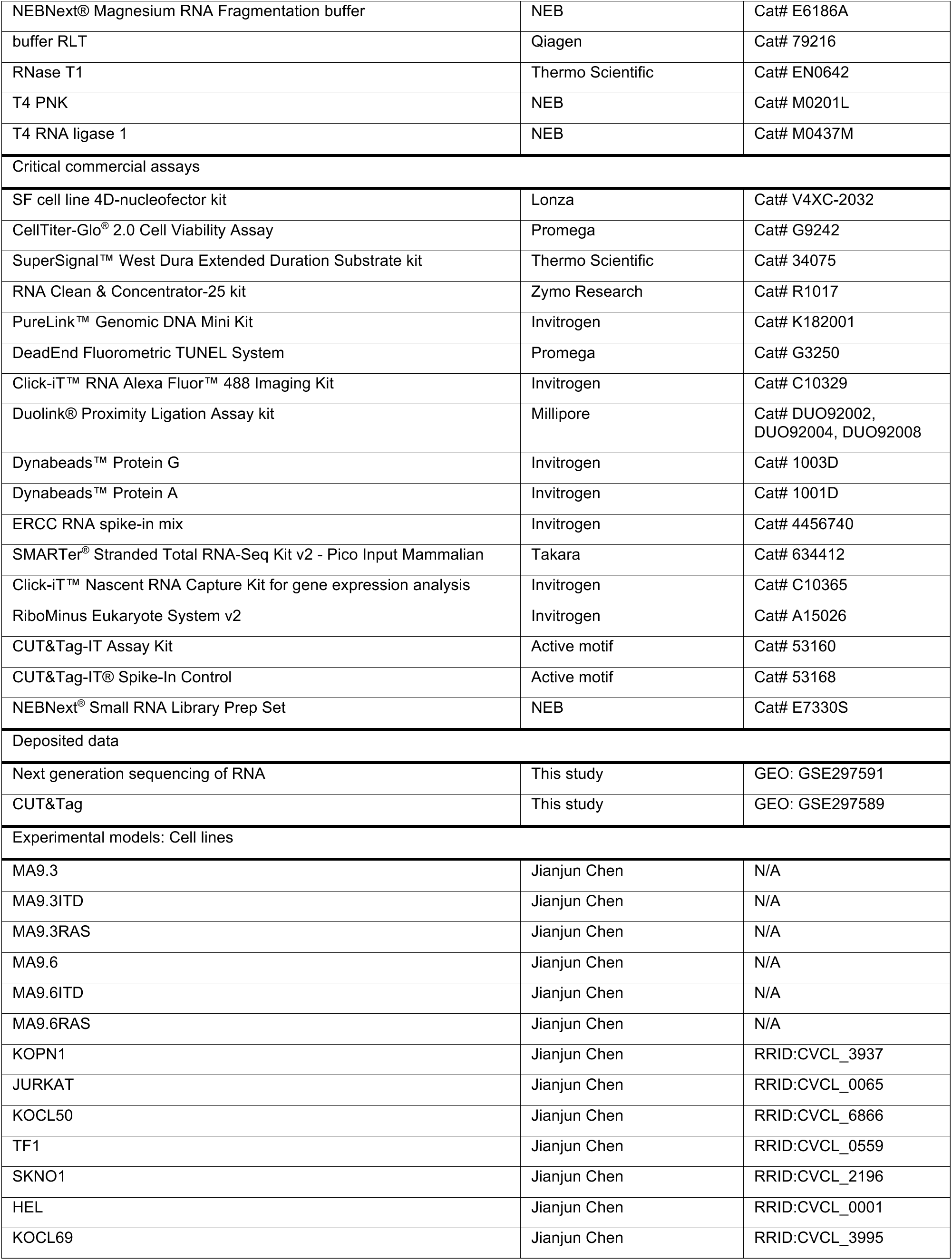
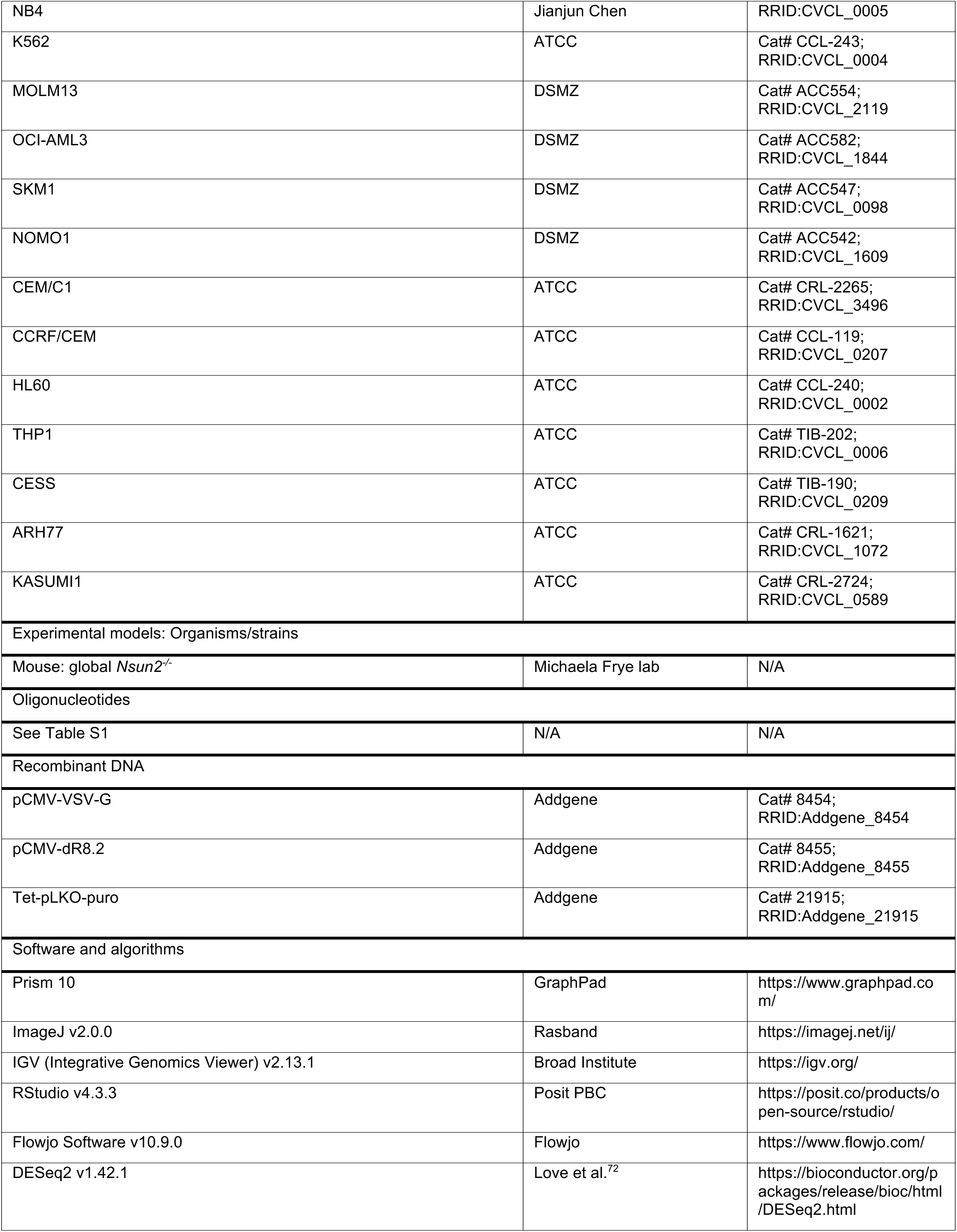
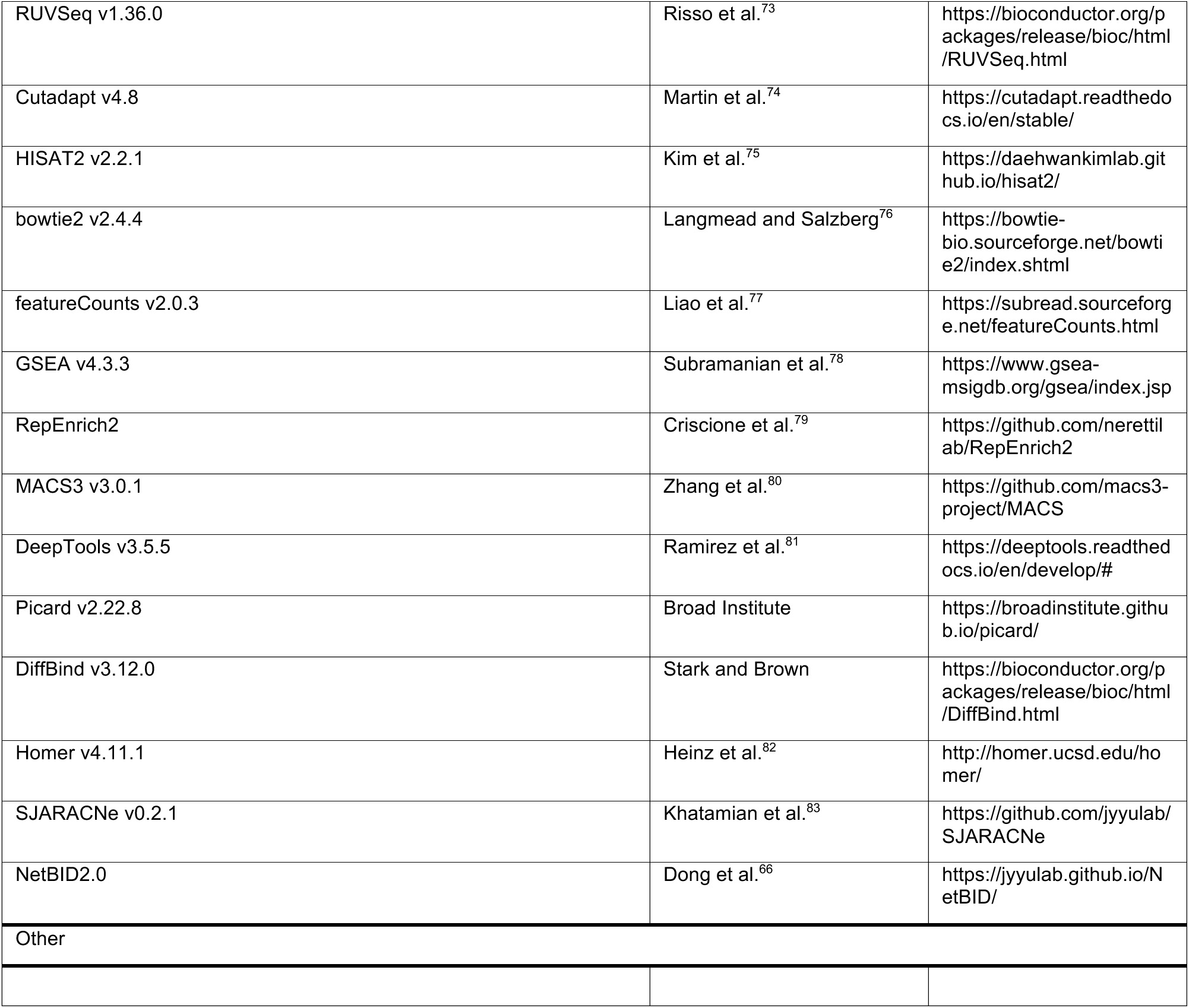

### METHOD DETAILS

#### Cell culture and cell line construction

The following cell lines were kindly provided by the J.Chen laboratory^84^: MA9.3, MA9.3ITD, MA9.3RAS, MA9.6, MA9.6ITD, MA9.6RAS, KOPN1, SKNO1, KOCL69, JURKAT, KOCL50, TF1, HEL, and NB4.

K562, MOLM13, OCI-AML3, SKM1, NOMO1, CEM/C1, CCRF/CEM, HL60, THP1, CESS, ARH77, KOPN1, KOCL69, JURKAT, KOCL50, HEL, and NB4 were maintained in RPMI1640 medium supplemented with 10% FBS and 50 U/mL Penicillin-Streptomycin. KASUMI1 cells were cultured in RPMI1640 with 10% FBS and 50 U/mL P/S. TF1 cells were kept in RPMI1640 with 10% FBS, 50 U/mL P/S and 2 ng/mL human GM-CSF. SKNO1 cells were grown in RPMI1640 with 10% FBS, 50 U/mL P/S and 10 ng/mL human GM-CSF. MA9.3 and MA9.6 cells were cultured in IMDM supplemented with 20% FBS, 50 U/mL P/S, 10 ng/mL each of human cytokines SCF, TPO, FLT3L, IL-3, and IL-6. MA9.3ITD, MA9.3RAS, MA9.6ITD and MA9.6RAS were kept in IMDM with 20% FBS and 50 U/mL P/S. Mycoplasma contamination was routinely tested using qPCR every half a year.

Lentiviral transduction was performed using viral particles packaged with HEK293T cells with pCMV-VSV-G and pCMV-dR8.2 plasmids (Addgene 8454 and 8455). Viral particles were precipitated using TransDux MAX Lentivirus Transduction Reagent and used immediately following overnight precipitation.

Transfection of K562 cells was conducted using Lipofectamine™ LTX Reagent with PLUS™ Reagent, following the manufacturer’s recommended protocol for K562. Cells were cultured to reach a concentration of 0.8 million/mL, and suspended in 25 mL Opti-MEM™ I Reduced Serum Medium. A DNA mixture containing 30 μg plasmid and 30 μL PLUS™ Reagent in 3.75 mL Opti-MEM™ medium was incubated at room temperature (RT) for 10 mins. Subsequently, 120 μL LTX Reagent was added and incubated for 25 mins at RT before being introduced to the cell culture. Transfection was conducted for 8 hours before replacing the media with standard growth medium.

Nucleofection of HEL cells was performed using Lonza 4D-Nucleofector System with the SF cell line 4D-nucleofector kit according to the manufacturer’s protocol. Each reaction utilized 1 million cells freshly passaged at 0.2 million/mL the day before. Cells were electroporated using program DC102.

#### Cell proliferation assay and 5-azaC sensitivity screen

Cell lines expressing Dox-induced shRNA were treated with the indicated compounds following 5 days of Dox induction with replenishment every 24 hours. As an exception, proliferation of MOLM13 with NSUN2 depletion was measured from the first treatment of Dox. In continuous measurement of proliferation, cells were passaged every 2 days, and the cumulative proliferation was adjusted by accounting for passaging ratios during proliferation measurement. HEL cells subjected to gene overexpression were treated with drugs 24 hours post-nucleofection. All proliferation assays were initiated at a seeding density of 0.2 million/mL and maintained with drug treatments for the specified time course. Cell proliferation was quantified using CellTiter-Glo^®^ 2.0 Cell Viability Assay, according to the manufacturer’s protocol with modifications for enhanced lysis efficiency. Specifically, CellTiter-Glo^®^ 2.0 reagent was freshly diluted in RIPA buffer (25 mM Tris-HCl pH 7.6, 150 mM NaCl, 1% NP-40, 1% Na-deoxycholate, 0.1% SDS) at a 3:4 volume ratio. For each measurement, 100 μL of diluted reagent was added to 100 μL of cell culture, mixed vigorously for 2 mins using orbital shaking in “fast” mode, and incubated at RT for 10 mins. Luminescence was recorded using a BioTek Synergy HTX plate reader.

Proliferation responses to 5-azaC and decitabine across 26 leukemi cells lines were evaluated after 24-hour treatment across five concentrations. Proliferation profiles were clustered based on normalized luminescence signals across doses. The top 50 recurrent leukemia-associated mutations were selected for visualization, based on the number of mutated cases in leukemia reported by the curated set of non-redundant studies reported in cBioPortal^85–87^. Mutation annotations were retrieved from the DepMap database^67^ and supplemented using data from COSMIC^88^ and Cellosaurus^89^. Silent mutations were excluded from the final integrated mutation dataset.

### Human primary AML patient samples

Human primary AML patient samples were collected at the time of diagnosis, relapse or remission after written informed consent at City of Hope Hospital or MD Anderson Cancer Center incongruence with the protocol approved by the institutional review board (IRB). Characteristics of the AML patients are provided in Table S3. The AML patient samples were cryopreserved in liquid nitrogen for further study. AML #2017-129, #2016-35 and #6303 cells were cultured in StemSpan^TM^ SFEMII supplemented with 10ng/mL of human SCF, mouse TPO, mouse FLT3L, human IL-3, human IL-6, 35 nM of UM171, 500 nM of Stem regenin1 (SR1) and 50 U/mL of P/S.

#### Animals

The global *Nsun2* KO mouse model was obtained from Frye’s lab^90^. These mice used in this study were backcrossed for more than 6 generations with C57BL/6 mice and applied throughout this study, including both male and female. All animal studies were performed with the approval from the Institutional Animal Care and Use Committee (IACUC) at The University of Texas Health Science Center at San Antonio (UTHSCSA) and conducted in accordance with the institutional and national guidelines and regulations.

#### *In vitro* colony-forming assay

Bone marrow (BM) cells were collected from wild-type or *Nsun2* KO mice and plated in triplicate in methylcellulose medium supplemented with 100 ng/mL mouse SCF, 10 ng/mL mouse IL-3, 50 ng/mL mouse TPO, 10 ng/mL mouse GM-CSF, 4 U/mL human EPO, and 50 ng/mL human interleukin-6. The colonies were imaged by STEMvision™ (STEMCELL Technologies) and scored on day 7. Colony cells were harvested and analyzed for expression of stem and progenitor markers and erythroid linage markers by flow cytometry.

#### Flow cytometry analysis

Flow cytometric analysis of mouse BM cells was performed as previously described^91^. Briefly, for HSPC cells, BM cells were stained with APC-anti-Lineage cocktail, PE-Cy^TM^7-anti-Sca1, PE-anti-c-kit, Alexa Flour 700-anti-CD34, APC-Cy7-anti-CD16/32, BV421-anti-CD135. For analysis of myeloid lineage cells, BM cells were stained with PerCP-Cy^TM^5.5-anti-Gr-1 and PE-anti-CD11b (Mac-1). APC-anti-Ter119 and PE-anti-CD71 were used for analyzing erythroid populations in BM or spleen cells.

#### Immunoblotting

Samples were collected with equal cell numbers and washed with PBS. For histone analysis, cells were lysed directly in 50 μL of 2x NuPAGE™ LDS sample buffer supplemented with β-mercaptoethanol (β-ME) per 1 million cells. To measure γH2A.X levels after UV damage, cells were exposed too 50 J/m^2^ UVC in PBS, then immediately incubated in complete medium for 5 mins prior to collection for immunoblotting.

For analysis of non-chromatin-associated proteins, cells were lysed in 50 μL of RIPA buffer supplemented with 100x proteinase inhibitor cocktail per 1 million cells on ice for 15 mins. Lysates were then centrifuged at 12,000 x g for 15 mins at 4°C. The supernatant was mixed with NuPAGE™ LDS sample buffer containing β-ME to a final 1x concentration. Samples were denatured by heating at 95°C for 20 mins and sonicated until no longer viscous. The denatured protein samples were separated on 4–12% NuPAGE Bis-Tris gels (Invitrogen™, NP0335BOX) and transferred to 0.45 μm nitrocellulose membrane (Bio-Rad, 1620115) using a semi-dry transfer system at 25 V for 45 mins. Membranes were blocked in 5% BSA in PBSTat RT for 30 mins, followed by overnight incubation at 4°C with primary antibody diluted in PBST containing 3% BSA. After three 10-min washes in PBST, membranes were incubated with secondary antibody diluted in PBST containing 1% BSA for 40 mins at RT, washed three more times for 10 mins each, and developed using SuperSignal™ West Dura Extended Duration Substrate kit. Signals were visualized with the Invitrogen™ iBright imaging system. The SRSF2 antibody used for immunoblotting is Cat# ab204916 (Abcam).

#### Cellular fractionation and purification of caRNA

Cellular fractionation was performed based on previously described protocol^38^ with slight modifications. A total of 20 million cells were washed with PBS and pelleted by centrifugation at 500 x g. Cells were resuspended in 300 μL of ice-cold NP-40 lysis buffer (10 mM Tris-HCl pH 7.5, 150 mM NaCl, 0.05% NP40) by gently flicking the tube, avoiding pipetting. After 5 mins of incubation on ice, the suspension was layered onto ice-cold NP-40 lysis buffer containing 24% sucrose and centrifuged at 15,000 x g for 10 mins at 4°C. The supernatant was collected as the cytoplasmic fraction. The pellet was washed with ice-cold PBS and resuspended in 150 μL of ice-cold glycerol buffer (20 mM Tris-HCl pH 8.0, 75 mM NaCl, 0.5 mM EDTA, 50% glycerol) by flicking the tube. Next, 300 μL of ice-cold nuclei lysis buffer (10 mM HEPES pH 7.6, 7.5 mM MgCl2, 0.2 mM EDTA, 300 mM NaCl, 1M Urea, 1% NP40) was added to the suspension, which was then vortexed at maximum speed for 20 seconds, incubated on ice for 2 mins, and centrifuged at 15,000 x g for 2 mins at 4°C. The supernatant was collected as the nuclear fraction.

The remaining pellet was washed with PBS and harvested as the chromatin fraction.

To isolate caRNA, the chromatin pellet was treated with 2 U TURBO DNase™ at 37 °C for 1 hour, followed by digestion with proteinase K at 55°C for 1 hour after supplementing 1% SDS to the buffer. TRIzol™ reagent was added to the mixture, which was then incubated at 50°C for 20 mins for resolution of pellets. Following TRIzol™ extraction with chloroform, the aqueous phase was collected, and RNA was purified using the RNA Clean & Concentrator-25 kit (RCC-25) including in-column DNase I digestion according to the manufacturer’s instructions.

#### UHPLC–MS/MS

70 ng of RNA was digested with nuclease P1 at 37 °C overnight, followed by treatment with 1 U shrimp alkaline phosphatase (rSAP) at 37 °C for 2 hours. Genomic DNA, purified from PureLink™ Genomic DNA Mini Kit, was digested with nuclease P1 at 37 °C overnight. This was followed by sequential digestion with 0.001 U phosphodiesterase I in 100 mM ammonium bicarbonate at 37 °C for 2 hours, and 1 U rSAP at 37 °C for 2 additional hours. Prior to analysis, all samples were diluted with LC-grade water and filtered through a 0.22 μm PVDF filter (Millipore, SLGVR04NL) prior to injection into a C18 reversed-phase column connected to a SCIEX 6500+ Triple Quadrupole Mass Spectrometer operating in positive electrospray ionization mode. Nucleosides were quantified based on retention time and specific nucleoside-to-base ion transitions (RNA: 268 → 136 for A; 258 → 126 for m^5^C; DNA: 252 → 136 for dA, 242 → 126 for 5mdC). Relative nucleoside levels were determined by peak area comparison within the same batch of reactions.

#### TUNEL assay

TUNEL assay was performed using the DeadEnd Fluorometric TUNEL System according to the manufacturer’s instructions, with the modification that the PBS wash buffer was supplemented with 0.5% BSA. Following rTdT labeling, the reaction was terminated using 20 mM EDTA containing 0.5% BSA in PBS to prevent cell clumping.

#### EU-based nascent RNA labeling analysis

Nascent RNA labeling for fluorescence detection was performed using the Click-iT™ RNA Alexa Fluor™ 488 Imaging Kit. Cells were incubated with 1 mM EU for 30 mins prior to harvesting. The procedure followed the manufacturer’s instructions, with modifications for flow cytometry analysis. At each wash step, cells were washed using 0.5% BSA in PBS to minimize cell clumping, and centrifuged at 3,000 x g for 3 mins at RT-except during the rinsing step, where the Click-iT^®^ reaction rinse buffer was used as instructed.

#### EdU-based cell cycle analysis and unscheduled DNA repair analysis

For cell cycle analysis, cells were incubated with 20 μM EdU for 1 hour, followed by a 30-min chase in complete media supplemented with 10 μM thymidine to dilute unincorporated EdU. Unscheduled DNA repair analysis was performed based on a published protocol^92^ with modifications. After treatment under designated conditions, cells were irradiated with 50 J/m^2^ 254 nm UVC in PBS, followed by immediate addition of complete media containing 20 μM EdU and 1 μM 5-Fluoro-2’-deoxyuridine. Cells were incubated for 1 hour under standard culture conditions, then washed twice with PBS and chased in complete media supplemented with 10 μM thymidine for 30 mins. At harvest, EdU detection was performed using Click-iT™ RNA Alexa Fluor™ 488 Imaging Kit, following a flow cytometry-adapted protocol as described for EU-based nascent RNA labeling. Post click chemistry, cells were stained with 2 μg/mL Hoechst 33342 for 15 mins at RT, washed, and analyzed by flow cytometry. Hoechst signal was quantified in linear scale. Cells with increased EdU incorporation and intermediate Hoechst intensity were classified as S phase, while 1N and 2N populations with basal level EdU intensity were designated as G0/G1 and G2/M phases, respectively. For unscheduled DNA repair analysis, only G0/G1 or G2/M populations were included in the quantification.

#### Immunofluorescence staining

Chamber slides were pre-coated with 0.1 mg/mL poly-D-lysine overnight to promote cell adherence. After harvest, cells were fixed with 4% paraformaldehyde (PFA) in PBS at RT for 10 mins, washed, and resuspended in PBS at a concentration of 1 million cells/mL. Cells were added to the pre-treated chambers and allowed to settle by gravity for 30 mins at RT without disturbance. Following sedimentation, excess PBS was gently removed, and cells were permeabilized by 0.5% Triton X-100 in PBS for 10 min, followed by PBS washes at RT. Cells were then blocked with 0.5% BSA in PBS or 20 min, and subsequently incubated with primary antibodies diluted in blocking buffer: anti-SRSF2 (ab11826) at ratio of 1:200 and anti-p300 at ratio of 1:400 for 1 hour at RT. After PBS washing, cells were incubated with secondary antibodies diluted 1:1000 in blocking buffer for 1 hour: goat anti-mouse IgG-AF 546 and goat anti-rabbit IgG-AF 647. Cells were then stained with 2 μg/mL Hoechst 33342 in PBS for 15 mins at RT. Following final PBS washes, ProLong™ Glass Antifade Mountant was applied to minimize photobleaching. Fluorescence images were acquired using a Leica SP8 laser scanning confocal microscope. The fluorescence intensity was measured by ImageJ.

#### Proximity ligation assay

Proximity ligation assay was conducted using the Duolink® Proximity Ligation Assay kit. Cells were fixed with 4% PFA in PBS at RT for 10 mins, followed by gravity sedimentation at a concentration of 1 million cells/mL onto chamber slides pre-coated with 0.1 mg/mL poly-D-lysine overnight. After permeabilization with 0.5% Triton X-100 in PBS for 10 min, the assay was carried out according to the manufacturer’s protocol. Primary antibodies used were anti-SRSF2 (ab11826, 1:200) and anti-p300 (1:400). Following amplification, nuclei were stained with 2 μg/mL Hoechst 33342 for 15 mins, and slides were mounted using ProLong™ Glass Antifade Mountant. Fluorescence imaging was conducted using a Leica SP8 laser scanning confocal microscope.

#### Immunoprecipitation

Immunoprecipitation was performed using 50 μL of Dynabeads™ Protein G or Dynabeads™ Protein A, washed with lysis buffer (50 mM Tris-HCl pH 7.5, 100 mM NaCl, 1% NP-40 substitute, 0.5% sodium deoxycholate, 100 × proteinase inhibitor cocktail). Beads were conjugated with SRSF2 antibodies-20 μL of the one from Abcam (ab204916), 5 μg of the one from Proteintech or 5 μL of the one from Sigma-Aldrich-and Rabbit IgG or Mouse (G3A1) mAb IgG1 Isotype Control as controls. Protein G beads were used for mouse antibodies and protein A beads were used for rabbit antibodies. 20 million SKM1 cells per sample were lysed in 800 μL lysis buffer at 4 °C with end-to-end rotation for 30 mins. Lysates were centrifuged at 15,000 x g for 15 min at 4 °C, and the supernatant was collected. 40 μL of lysate was reserved as input, and the remainder was incubated with the antibody-conjugated beads at 4°C for 4 hours. Beads were washed three times with wash buffer (50 mM Tris-HCl pH 7.5, 300 mM KCl, 0.05% NP-40 substitute, 1000 × proteinase inhibitor cocktail). washed beads were suspended in 50 μL of 1x Laemmli sample buffer supplemented with β-ME and heated at 95 °C for 10 minutes. The input sample was mixed with 4x Laemmli sample buffer supplemented with β-ME to a final 1x concentration and heated the same. 5 μL of input and 20 μL of eluted sample were loaded for immunoblotting.

#### RNA-seq

RNA extraction was performed using TRIzol™ reagent following the designated treatment. The RNA was extracted with chloroform, and the aqueous phase was retained for RNA purification using RCC-25. In-column DNase I digestion was conducted to remove DNA. For studies involving gene depletion, KD efficiency was assessed by RT-qPCR. For RNA-seq after 5-azaC treatment, 2 μL of a 1/20,000 dilution of ERCC RNA spike-in mix was added to 10 ng of RNA. The RNA was then proceeded for library construction using the SMARTer^®^ Stranded Total RNA-Seq Kit v2 - Pico Input Mammalian. Next-generation sequencing was performed on Illumina NovaSeqX with pair-end 150 bp reads.

#### Nascent RNA-seq

MOLM13 and SKM1 cells were treated with 3 μM 5-azaC for 0, 4, 8 and 16 hours. Following treatment, 1 mM EU was added to the media for nascent RNA labeling. In parallel, *drosophila* cells were labeled similarly to serve as a spike-in control. Cells were harvested and purified using TRIzol™ reagent following the manufacturer’s protocol with isopropanol precipitation. Nascent RNA was enriched using the Click- iT™ Nascent RNA Capture Kit for gene expression analysis, following the manufacturer’s protocol with modifications. 4.5 μg of EU-labeled RNA was reacted with 2.2 μL biotin azide in a 25 μL click reaction and purified using RCC-25. 600 ng of RNA was mixed with *drosophila* spike-in at a 200:1 ratio and processed for nascent RNA enrichment. Streptavidin beads provided by the kit were washed with B&W buffer (5 mM Tris-HCl pH 7.5, 0.5 mM EDTA, 1M NaCl) three times, followed by solution A (100 mM NaOH, 50 mM NaCl) twice for 2 mins each. The beads were then washed with 100 mM NaCl at the same volume of solution A, followed by another round of washing with B&W buffer three times. Beads were suspended in the wash buffer 2 provided by the kit, and RNA was mixed with RNaseOUT in RNA binding buffer and denatured at 70°C for 5 mins before being added to the beads. After 30 mins of incubation at RT with end-to-end rotation, the beads were washed with low salt buffer (10 mM Tris-HCl pH 7.5, 50 mM NaCl, 0.1% NP-40, 0.2 U/μL SUPERase•In™ RNase Inhibitor), high salt buffer (10 mM Tris-HCl pH 7.5, 500 mM NaCl, 0.1% NP-40, 0.2 U/μL SUPERase•In™ RNase Inhibitor), and B&W buffer supplemented with 0.2 U/μL SUPERase•In™ RNase Inhibitor, washing five times each. The beads were then washed with the wash buffer 1 and wash buffer 2 supplied by the kit and eluted in 50 μL of 95% formamide supplemented with 2.5 mM EDTA by heating at 65°C for 5 mins, followed by 95°C for 5 mins. The supernatant was immediately harvested while still hot, and RNA was extracted using 1 mL TRIzol™ reagent. RNA was purified following the manufacturer’s protocol with isopropanol precipitation facilitated by glycogen. The library was constructed using the SMARTer^®^ Stranded Total RNA-Seq Kit v2 - Pico Input Mammalian, and sequencing was performed using the Illumina NovaSeqX with pair-end 50 bp reads.

#### m^5^C MeRIP-seq

m^5^C MeRIP-seq was performed as described in previous publication^30^. For MOLM13 cells, 0.02 million *drosophila* S2 cells were added to 20 million MOLM13 cells before proceeding with cellular fractionation, as outlined earlier. rRNA removal was carried out using RiboMinus Eukaryote System v2. 2 μg ribo-depleted caRNA was then used for m^5^C immunoprecipitation. RNA was fragmented using 1× NEBNext® Magnesium RNA Fragmentation buffer at 94°C for 2 mins, and 5% of the sample was saved as input. The RNA was heated to 72°C for 2 mins and then placed immediately on ice for denaturation. Next, the RNA was added to protein G dynabeads conjugated with 2 μg anti-m^5^C antibody in 300 μL IP buffer (10 mM Tris-HCl pH 7.5, 150 mM NaCl, 0.05% Triton X-100, 1 mM spermidine, 0.2 U/μL SUPERase•In™ RNase Inhibitor), and incubated with end-to-end rotation at 4°C for 2 hours. The beads were washed three times with IP buffer and eluted with buffer RLT. Both input and MeRIP libraries were constructed using the SMARTer^®^ Stranded Total RNA-Seq Kit v2 - Pico Input Mammalian, and sequencing was performed with the Illumina NovaSeqX with pair-end 150 bp reads.

#### CUT&Tag

CUT&Tag was performed using the CUT&Tag-IT Assay Kit following the manufacturer’s instructions. 0.2 million cells were used per sample. For 5-azaC-treated CUT&Tag profiling of H2AK119ub and H3K27ac, 4,000 *drosophila* nuclei from CUT&Tag-IT® Spike-In Control were added as a spike-in control. Primary antibodies used included 1 μL H2AK119ub antibody, 1 μL H3K27ac antibody (Cat# 39034) or 0.25 μL SRSF2 antibody (Cat# ab11826). Libraries were constructed according to the kit protocol and sequenced on the Illumina NovaSeqX platform with pair-end 150 bp reads.

#### CLIP-seq and CLIP with end ligation of pCp-AF488 fluorophore

For CLIP-seq, 100 million cells were used for each replicate. Following the designated treatment, cells were crosslinked in PBS using 4500 J/m^2^ UV at 254 nm. Cells were lysed in 200 μL iCLIP lysis buffer (50 mM Tris-HCl pH 7.5, 100 mM NaCl, 1% NP-40 substitute, 0.1% SDS, 0.5% sodium deoxycholate, 100× proteinase inhibitor cocktail and 0.2 U/μL SUPERase•In™ RNase Inhibitor) at 4°C for 15 mins with end-to-end rotation. SDS were then added to reach a final concentration of 1% for protein release from chromatin. Samples were sonicated (35% amplitude, 2s on / 4s off cycles, 1 min on ice) and diluted 10- fold with SDS-free iCLIP lysis buffer to reduce SDS back to 0.1%. After centrifugation at 15,000 x g for 30 mins at 4°C, the supernatant was harvested and treated with 0.2 U/μL RNase T1 at 22°C for 15 mins, then quenched on ice for 5 mins. 2% of the lysate was saved as input and digested with 10 U/μL RNase T1 at 22°C for 10 mins, followed by proteinase K digestion, RNA purification and end repair. The remaining lysate was incubated overnight at 4°C with protein A beads conjugated to 30 μL MBD6 antibodies (gifted from Prof. Lu Wang^93^) per sample. Beads were washed three times with CLIP wash buffer (50 mM Tris-HCl pH 7.5, 300 mM KCl, 0.05% NP-40, 1000× protein inhibitor cocktail and 0.2 U/μL SUPERase•In™ RNase Inhibitor), followed by digestion with 10 U/μL RNase T1 at 22°C for 10 mins.

Beads were further washed for three times each with CLIP high salt buffer (50 mM Tris-HCl pH 7.5, 500 mM KCl, 0.05% NP-40, 100× protein inhibitor cocktail and 0.2 U/μL SUPERase•In™ RNase Inhibitor) and PNK buffer without DTT, and subjected to end-repair using T4 PNK. After proteinase K digestion, RNA was purified for library preparation. The input and CLIP libraries were constructed using the NEBNext^®^ Small RNA Library Prep Set and sequenced with Illumina NovaSeqX, pair-end 150 bp.

For CLIP with end ligation of pCp-AF488 fluorophore, K562 cells with SRSF2-myc overexpression after DMSO or 5-azaC treatment were counted to ensure the same number of cells used for experiment in each sample. Approximately 15 million cells were crosslinked by 4500 J/m^2^ UV at 254 nm in PBS. Samples were lysed with iCLIP lysis buffer, and lysates were cleared by centrifugation. Supernatants were treated with 0.2 U/μL RNase T1 at 22°C for 15 mins, quenched on ice, and incubated at 4°C for 4 hours with protein G beads conjugated to 4 μL Myc-tag antibodies. Following immunoprecipitation, beads were washed for three times with CLIP wash buffer, digested with 10 U/μL RNase T1 at 22°C for 8 mins, and further washed for three times each with CLIP high salt buffer and PNK buffer without DTT. Beads were subjected to RNA dephosphorylation using T4 PNK, then ligated overnight at 16°C in a 26 μL ligation reaction containing 2.4 μL T4 RNA ligase 1, 0.5 μL pCp-AF488, 0.3 μL 100 mM ATP, 0.4 μL SUPERase•In™ RNase Inhibitor, 0.6 μL 1% Tween-20, 0.9 μL DMSO and 9 μL 50% PEG8000. After ligation, samples were washed, denatured in 2x LDS sample buffer with β-ME at 70°C for 10 mins, and analyzed on a 4–12% NuPAGE Bis-Tris gel for fluorescence detection.

#### Data analysis of RNA-seq

Adapter trimming was performed using Cutadapt^74^, and reads were mapped to the hg38 human genome or mm10 mouse genome using HISAT2^75^. Gene-level read counts were quantified using featureCounts^77^ based on the hg38 refGene annotation. For differential gene expression analysis, DESeq2^72^ was applied to datasets without spike-in, while RUVSeq^73^ was used for normalization and analysis of datasets that included ERCC RNA spike-in mix. GSEA was conducted using the GSEA software suite^78,94^.

#### Data analysis of nascent RNA-seq

Adapter trimming was performed using Cutadapt^74^, and reads were mapped to hg38 human genome and dm6 *drosophila* genome by HISAT2^75^. Gene-level read counts were quantified using featureCounts^77^ based on the hg38 refGene annotation. For differential gene expression analysis, DESeq2^72^ was applied using the read counts mapped to *drosophila* chromosomes as spike-in normalization. GSEA was conducted using the GSEA software suite ^78,94^.

#### Data analysis of m^5^C MeRIP-seq

Adapter trimming was performed using Cutadapt^74^, and reads were deduplicated and aligned to the hg38 human genome and dm6 *drosophila* genome using HISAT2^75^. Gene-level read counts were quantified using featureCounts^77^ based on the hg38 refGene annotation. In parallel, reads mapping to repetitive elements were analyzed using Bowtie2^76^ followed by RepEnrich2^79^. input samples were analyzed as caRNA-seq for differential expression analysis of caRNA, covering both coding genes and repetitive elements. DESeq2^72^ was used for differential expression analysis. For samples from MOLM13 cells with *drosophila* spike-in, total read numbers mapped to each *drosophila* chromosome were used for normalization in DESeq2 analysis.

MeRIP peaks were identified using MACS3^80^, and read counts from both input and MeRIP libraries were quantified at peak regions to assess enrichment. Peaks were normalized to total mapped reads, or additionally normalized by *drosophila* spike-in for MOLM13 samples. Regions with MeRIP/input > 0 in either control or 5-azaC-treated samples were considered putative m^5^C sites. Sites with an enrichment fold change of 5-azaC control < 0.5 were defined as decreased m^5^C sites. Metaplots were generated using deepTools^81^. The IKZF1 ChIP data was downloaded from ENCODE experiment ENCSR948VFL.

#### Data analysis of CUT&Tag

Adapter trimming was performed using Cutadapt^74^, and reads were aligned to the hg38 human genome and dm6 *drosophila* genome using bowtie2^76^. Duplicate reads were removed using Picard MarkDuplicates^95^. For 5-azaC-treated histone modification profiling, signal normalization was performed based on the total number of reads mapped to *drosophila* genome. Peak calling was performed using MACS3^80^. For histone modifications, broad peaks were identified using the parameters “--broad --broad-cutoff 0.01” was used. Differential peak analysis was conducted using DiffBind^96^. For quantile-based analysis, peaks were annotated to the nearest RefSeq genes using Homer annotatePeaks.pl^82^, and assigned as their putative regulatory elements. Metaplots were generated with deepTools^81^.

#### Data analysis of CLIP-seq

Adapter trimming was performed using Cutadapt^74^, and reads were aligned to the hg38 human genome using HISAT2^75^. Duplicate reads were removed using Picard MarkDuplicates^95^. Metaplots were generated with deepTools^81^.

#### Inference of gene network

A gene regulatory network was inferred using SJARACNe^83^, based on gene expression profiles of all myeloid lineage leukemia cell lines available in the DepMap^67^ database. Candidate hub genes were selected from commonly mutated genes identified in MDS, AML and Myeloproliferative neoplasm, curated from non-redundant studies in cBioPortal^85–87^. Selection criteria included mutation frequency > 1%, mutation cases > 5 and total detected cases > 100. The network was further integrated with gene expression data from leukemia cell lines categorized as sensitive and resistant to 5-azaC, analyzed using NetBID2^66^.

### QUANTIFICATION AND STATISTICAL ANALYSIS

Data are plotted with error bars representing standard deviation. *P*-values indicated in the figures are unpaired, two-tailed, two-sided Student’s *t* test. An exception is the GSEA analysis, where the nominal *p-* values generated by the GSEA software are labeled in the figures.

## Supplemental information

**Figure S1.**
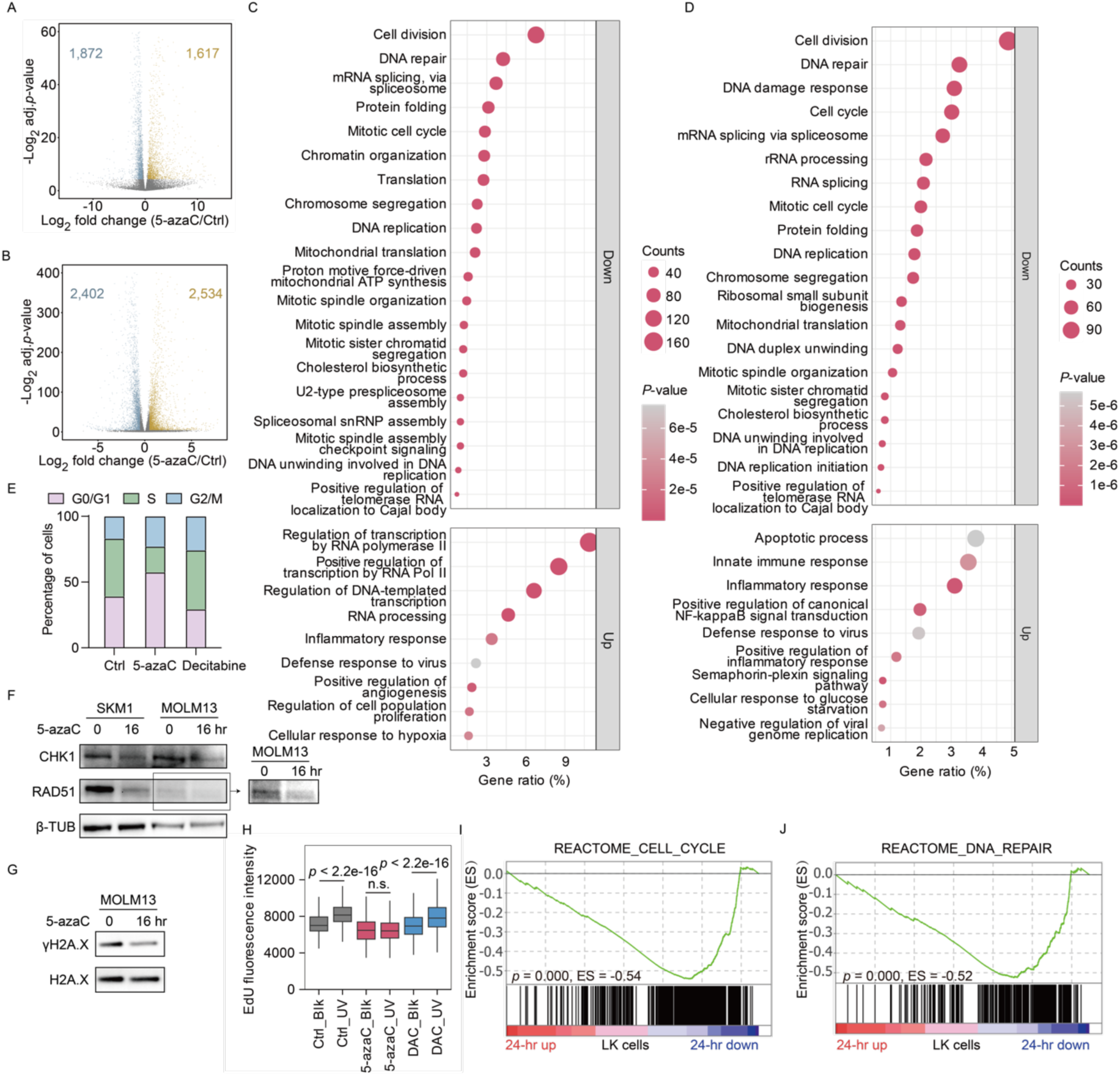
5-azaC modulates cell cycle, DNA damage repair, and apoptosis mediated by RNA m^5^C. (A and B) Volcano plot of RNA-seq from MOLM13 cells (A) and SKM1 cells (B) after a 24-hour 5-azaC (3 μM) treatment. DEGs are defined as |log2 fold change| > 0.58 and adj.*p*-value < 0.05. DEG numbers are annotated on the plot. Adj.*p*-value, adjusted *p*-value. (C and D) GO analysis of the top enriched biological processes in DEGs after 5-azaC (3 μM) treatment in MOLM13 (C) and SKM1 (D) cells. The top 20 GO terms enriched in downregulated genes and the top 9 GO terms enriched in upregulated genes with FDR < 0.05 were plotted. (E) Cell cycle distribution of SKM1 cells after 16 hours treatment with 3 μM of 5-azaC or decitabine. (F) Immunoblotting of DNA damage repair genes after 3 μM of 5-azaC treatment in MOLM13 cells. (G) Immunoblotting of γH2A.X after 3 μM of 5-azaC treatment followed by 50J/m^2^ UVC irradiation. (H) EdU incorporation during DNA damage repair after UVC irradiation in MOLM13 cells treated with 3 μM of 5-azaC or decitabine for 16 hours. Only G2/M-phase populations are shown. Blk, no irradiation. DAC, decitabine. (I and J) GSEA of MSigDB cell cycle (I) and DNA damage repair (J) gene set in 5-azaC-treated LK cells.

**Figure S2.**
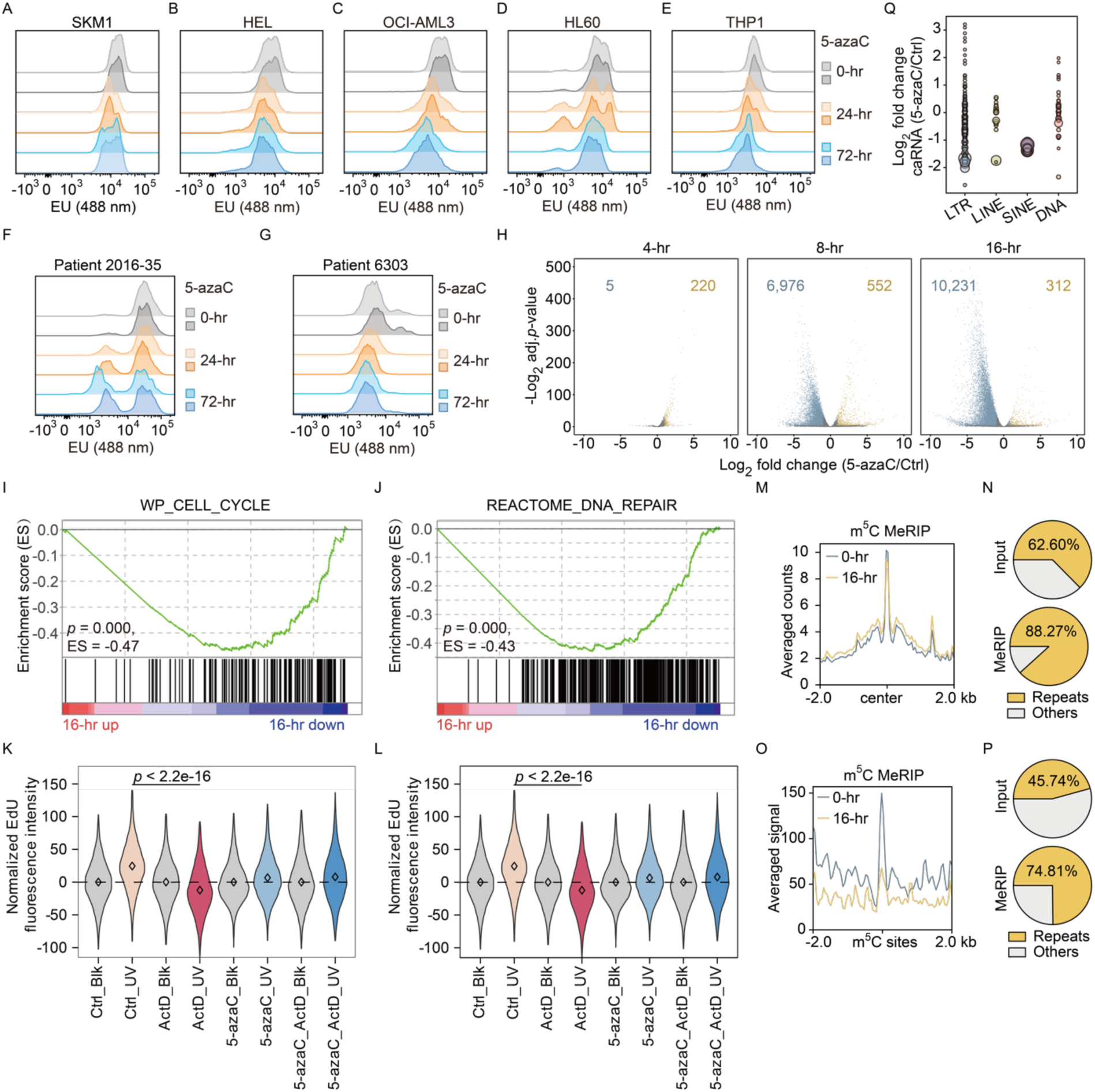
5-azaC induces transcriptional repression through caRNA m^5^C depletion. (A-G) EU fluorescent signals in SKM1 (A), HEL (B), OCI-AML3 (C), HL60 (D) and THP1 (E) cells and AML patient samples 2016-35 (F) and 6303 (G) after 3 μM of 5-azaC treatment. (H) Nascent RNA-seq of SKM1 cells after 3 μM of 5-azaC treatment. DEGs are defined as |log2 fold change (5-azaC/Ctrl) | > 1 and adj.*p*-value < 0.05. DEG numbers are annotated on the plot. Adj.*p*-value, adjusted *p*-value. (I and J) GSEA of transcriptional changes in cell cycle (I) and DNA damage repair (J) gene sets after 16 hours of 5-azaC (3 μM) treatment in SKM1. (K and L) EdU incorporation during DNA damage repair after UV irradiation in MOLM13 cells treated with 3 μM of 5-azaC for 16 hours. Samples annotated with ActD were treated with 1 μg/mL ActD for 2 hours prior to irradiation. Only G0/G1- (K) and G2/M- (L) phase populations are shown. Blk, no irradiation. (M) Metaplot of m^5^C MeRIP-seq signal changes at m^5^C sites from 5-azaC (3 μM) treated and non-treated MOLM13 cells. (N) Percentage of sequencing reads mapped to repetitive elements (Repeats) or not (Others) in input and MeRIP samples from control MOLM13 cells. The percentage is the average of three replicates. (O) Metaplot of m^5^C MeRIP-seq signal changes at m^5^C sites from 5-azaC (3 μM) treated and non-treated SKM1 cells. (P) Percentage of sequencing reads mapped to repetitive elements (Repeats) or not (Others) in input and MeRIP samples from control SKM1 cells. The percentage is the average of three replicates. (Q) Differential levels of caRNA repeat subfamilies in MOLM13 cells after a 16-hour 5-azaC (3 μM) treatment. Circle sizes represent normalized caRNA levels in untreated samples.

**Figure S3.**
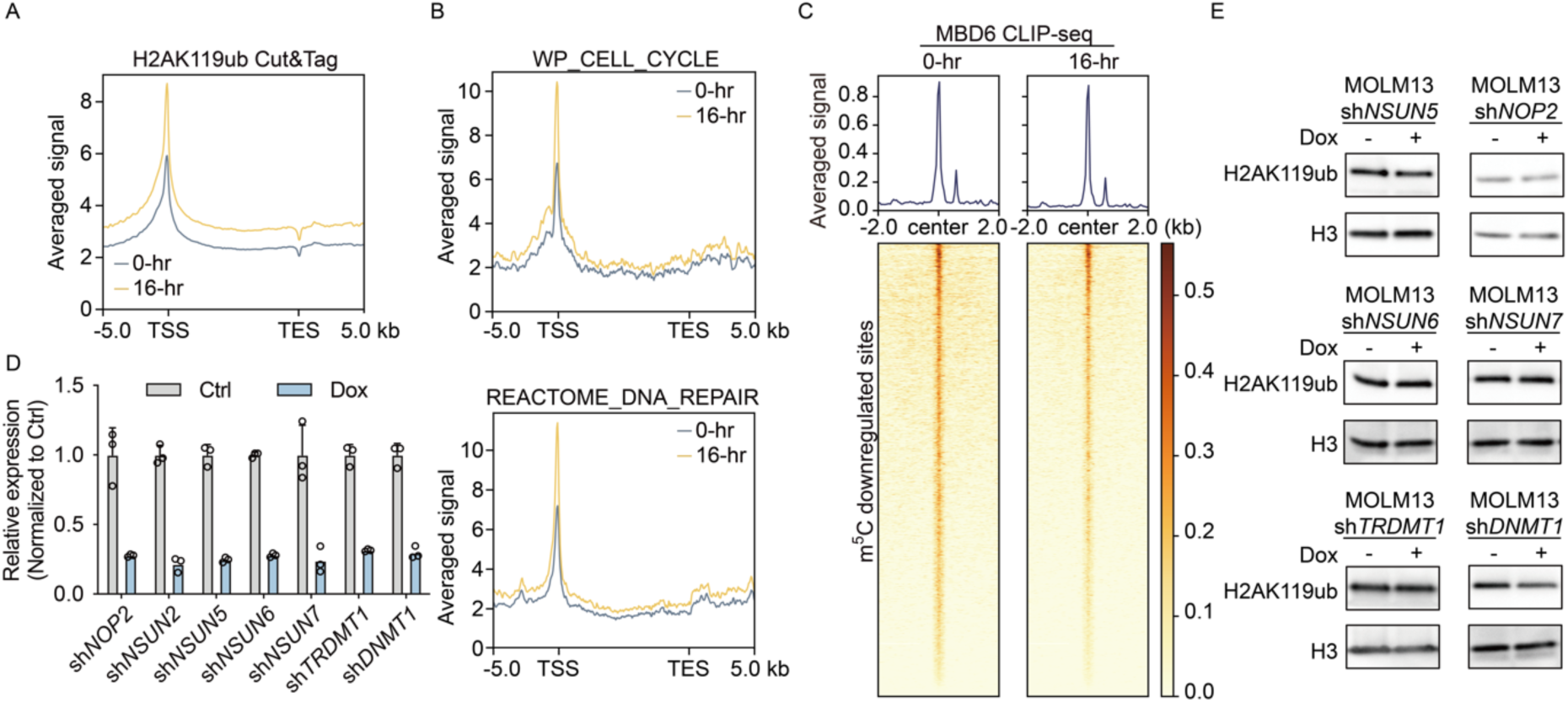
5-azaC induces transcriptional repression through m^5^C-dependent H2AK119ub regulation. (A and B) Metaplot of H2AK119ub changes across all annotated genes (A) or specific gene sets (B) after a 16-hour 5-azaC (3 μM) treatment in MOLM13 cells. (C) Metaplot and heatmap of MBD6 CLIP-seq signal changes at m^5^C sites that are downregulated after a 16-hour 5-azaC (3 μM) treatment in MOLM13 cells. (D) KD efficiency of m^5^C methyltransferases or *DNMT1* after a 5-day Dox-induced depletion of the respective gene. Expression levels were first normalized to *ACTB*, followed by normalization to the respective controls. (E) Immunoblotting of H2AK119ub changes following a Dox-induced depletion of m^5^C methyltransferases or *DNMT1* for 5 days in MOLM13 cells.

**Figure S4.**
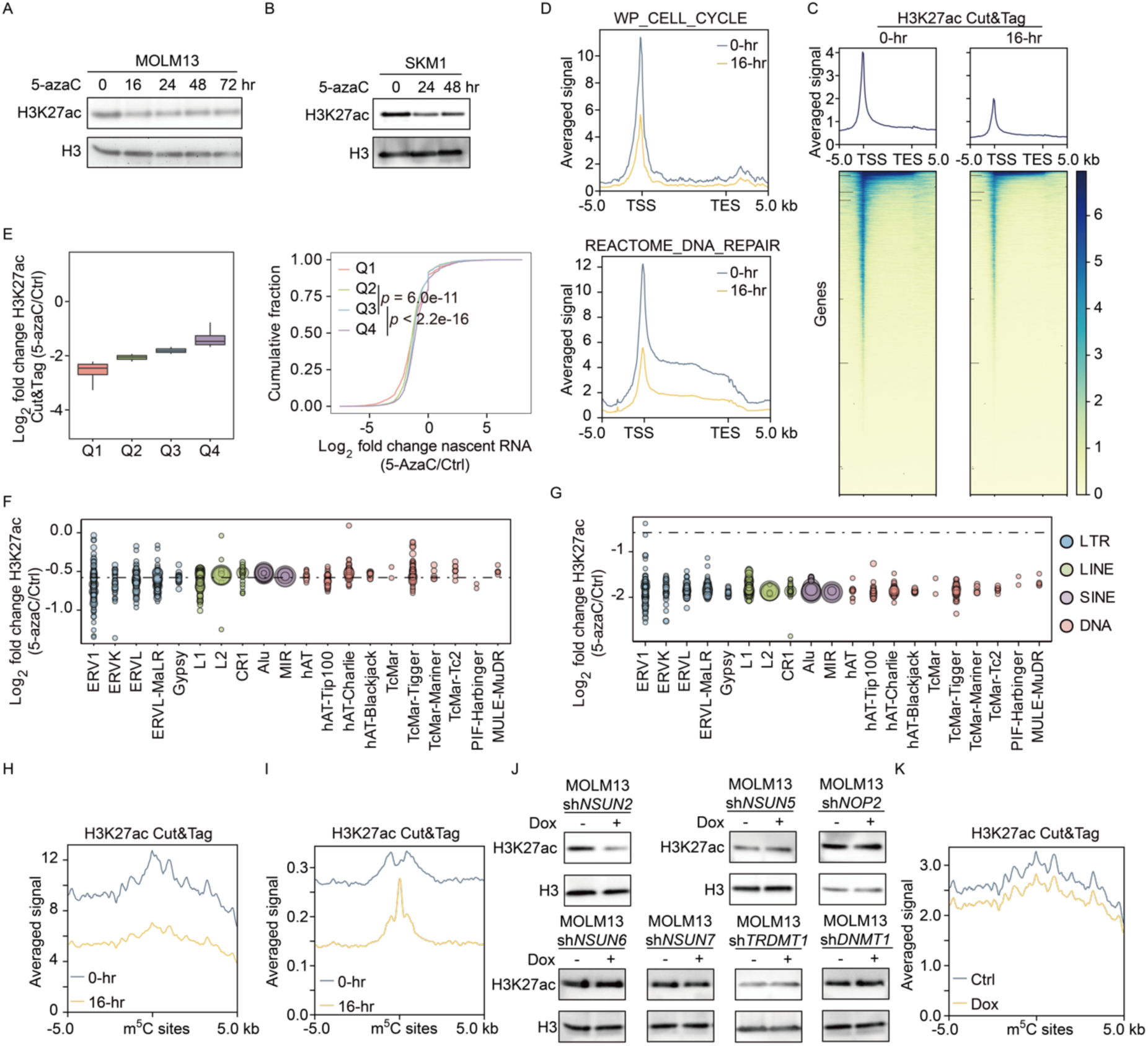
5-azaC induces transcriptional repression through H3K27ac downregulation. (A) Immunoblotting of H3K27ac changes in MOLM13 cells treated with 3 μM of 5-azaC. (B) Immunoblotting of H3K27ac changes in SKM1 cells treated with 1 μM of 5-azaC replenished every 24 hours. (C) Metaplot and heatmap of H3K27ac level changes across all annotated genes after a 16-hour 5-azaC (3 μM) treatment in MOLM13 cells. (D) Metaplot of H3K27ac changes in cell cycle (upper) and DNA damage repair (lower) gene sets in MOLM13 cells. (E) H3K27ac (left) and nascent RNA-seq (right) changes in genes grouped into four quantiles based on fold changes of nearby H3K27ac peaks in MOLM13 cells. Q1 represents the quantile with the most negative fold change. (F and G) H3K27ac changes across repeat families in SKM1 (F) and MOLM13 (G) cells. Circle sizes represent normalized CUT&Tag read counts in control samples. The dashed line indicates a value of -0.58. (H and I) Metaplot of H3K27ac level changes at m^5^C sites from the corresponding cell lines after 5-azaC (3 μM) treatment in SKM1 (H) and MOLM13 (I) cells. (J) Immunoblotting of H3K27ac changes following a Dox-induced depletion of RNA m^5^C methyltransferases or DNMT1 in MOLM13 cells. (K) Metaplot of H3K27ac level changes at m^5^C sites after a 5-day Dox-induced NSUN2 depletion in SKM1 cells.

**Figure S5.**
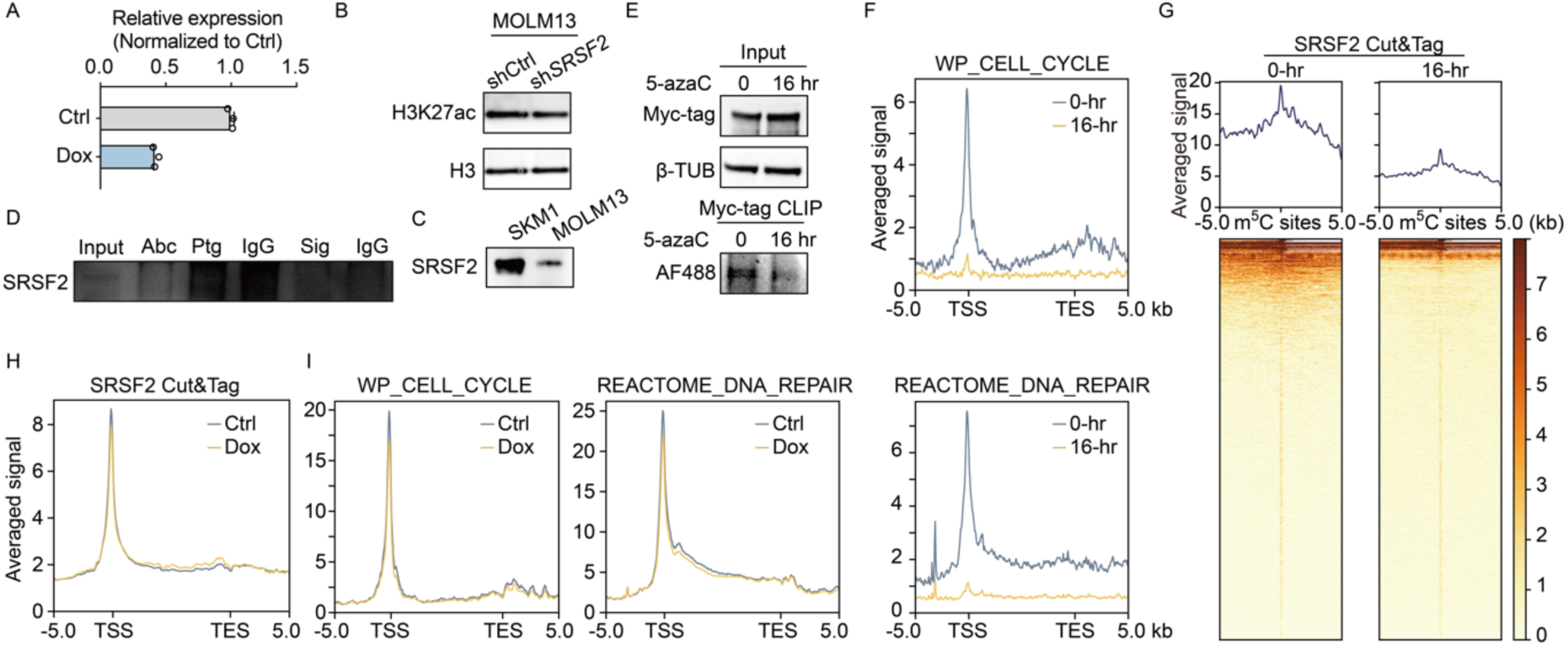
5-azaC downregulates H3K27ac through the SRSF2-mediated p300 recruitment. (A) KD efficiency of *SRSF2* after a 48-hour shRNA-mediated depletion. Expression levels were first normalized to *ACTB*, followed by normalization to the respective controls. (B) Immunoblotting of H3K27ac changes after a 48-hour shRNA-mediated SRSF2 depletion in MOLM13 cells. (C) Immunoblotting of SRSF2 levels across different cell lines. The samples were loaded with the same cell number. (D) Immunoprecipitation of SRSF2 with different antibodies: Abc (SRSF2 antibody ab204916), Ptg (SRSF2 antibody 20371-1-AP), Sig (SRSF2 antibody S4045). (E) Immunoblotting of SRSF2-Myc overexpression in K562 cells (upper) and Myc-tag CLIP followed by end ligation of pCp-AF488 fluorophore (lower). (F) Metaplot of SRSF2 CUT&Tag signal changes in cell cycle (upper) and DNA repair (lower) gene sets in SKM1 cells. (G) Metaplot and heatmap of SRSF2 CUT&Tag signal changes at m^5^C sites after a 16-hour 5-azaC (3 μM) treatment in SKM1 cells. (H and I) Metaplot of SRSF2 CUT&Tag signal changes across all annotated genes (H) or specific gene sets (I) following a 5-day Dox-induced NSUN2 depletion in SKM1 cells.

**Figure S6.**
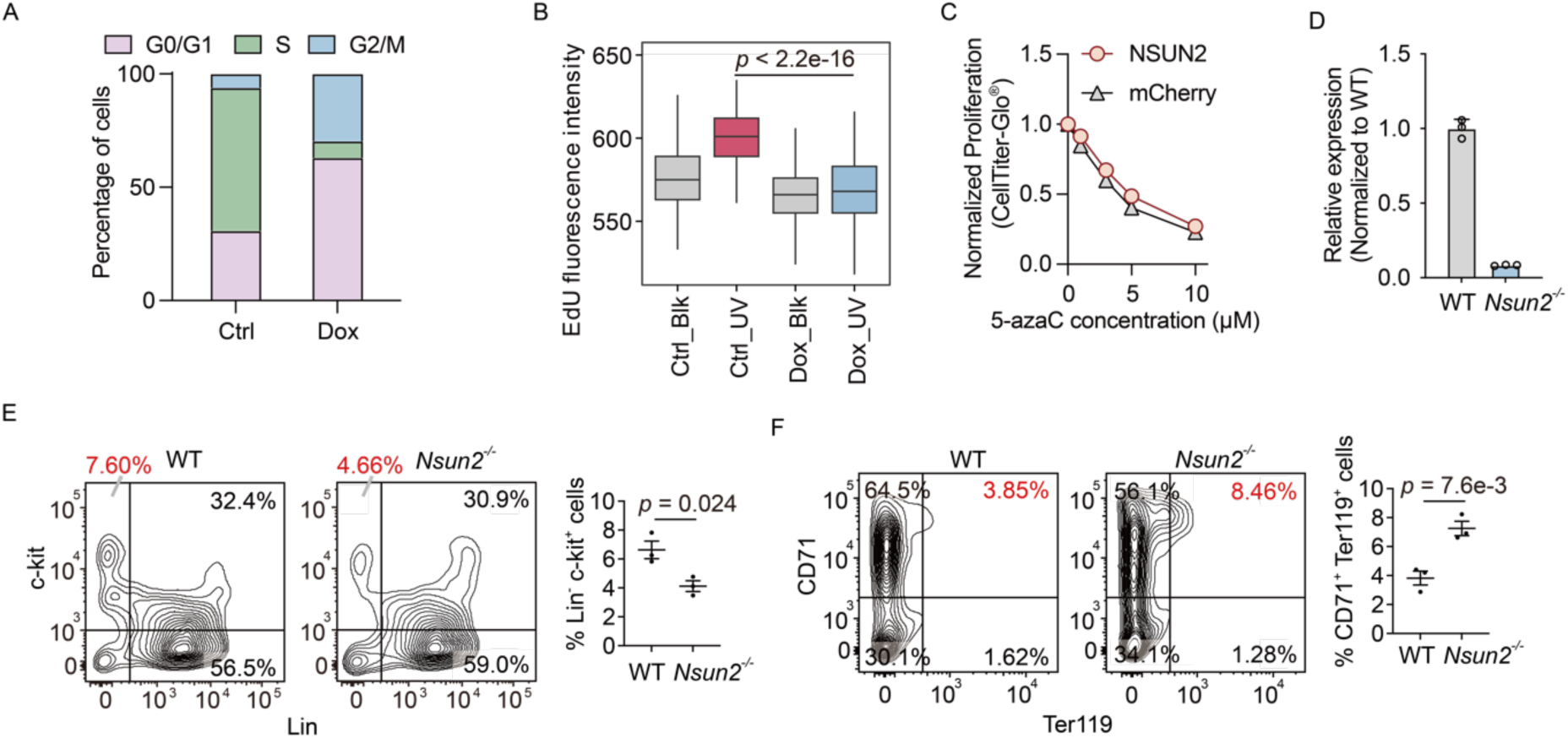
NSUN2 regulates leukemia proliferation and hematopoietic lineage commitment. (A) Cell cycle distribution of SKM1 cells after a 5-day Dox-induced NSUN2 depletion. (B) EdU incorporation during DNA damage repair after UVC irradiation in MOLM13 cells after a 5-day Dox-induced NSUN2 depletion. Only G2/M-phase populations are shown. Blk, no irradiation. (C) Cell proliferation of HEL with *NSUN2* overexpression followed by a 24-hour 5-azaC treatment. (D) KD efficiency of *Nsun2*^-/-^ mice measured in LK cells. (E and F) Flow cytometry analysis of LK (E) and erythroid (F) cell populations in progeny cells from WT and *Nsun2*^-/-^ CFU-C assays shown in Figure 6J.

**Figure S7.**
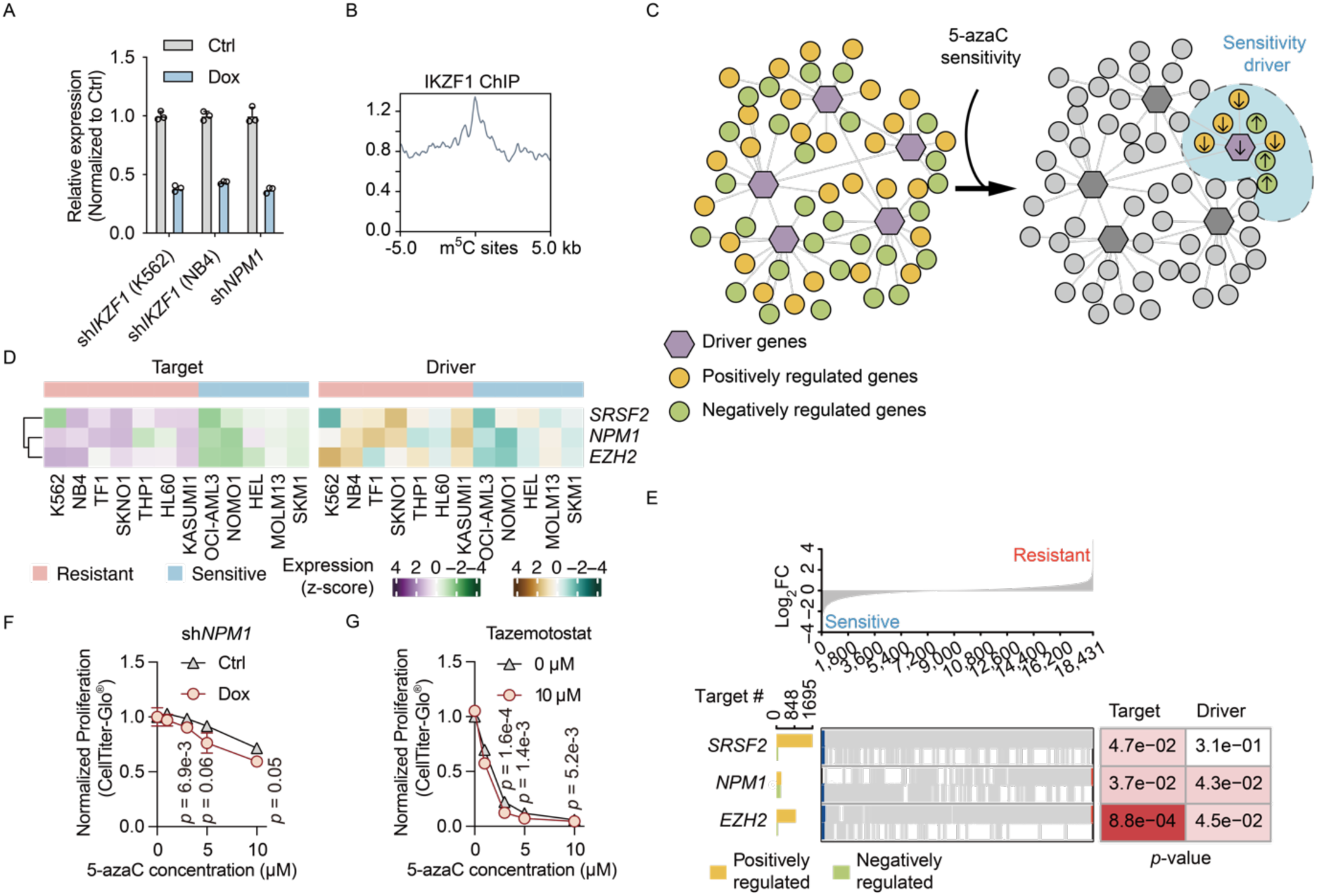
Gene network inference identifies mutations associated with 5-azaC sensitivity. (A) KD efficiency of *IKZF1* and *NPM1* after a 5-day Dox-induced depletion of the respective gene. Expression levels were first normalized to *ACTB*, followed by normalization to the respective controls. (B) Metaplot of IKZF1 ChIP-seq in K562 cells at m^5^C sites. (C) Schematic representation of the identification of genes determining 5-azaC sensitivity through gene network-based inference using NetBID2. (D) Expressions of frequently mutated genes in leukemia that correlate with 5-azaC sensitivity and their target genes in the network. (E) GSEA of positive and negative targets of frequently mutated genes in leukemia related to 5-azaC sensitivity. Log2FC refers to log2 fold change gene expression (resistant/sensitive). (F) Cell proliferation of THP1 following a 24-hour 5-azaC treatment after Dox-induced NPM1 depletion. (G) Cell proliferation of HEL following a 24-hour 5-azaC treatment after 48-hour 10 μM tazemotostat treatment.

